# Transient disruption of progenitor states and layer 2/3 projection neuron generation in frontal association cortex due to 22q11.2 gene deletion

**DOI:** 10.1101/2025.05.12.653556

**Authors:** Shah Rukh, Daniel W. Meechan, Connor Siggins, Zachary D. Erwin, Gia Baldo, Ashley Peck, Thomas M. Maynard, Anthony-Samuel LaMantia

**Affiliations:** Center for Neurobiology Research, The Fralin Biomedical Research Institute at Virginia Tech-Carilion School of Medicine, Roanoke, VA; Translational Biology, Medicine and Health Program; Department of Biological Sciences, Virginia Tech, Blacksburg, VA

## Abstract

The developmental origin of Layer 2/3 projection neuron (PN) pathology in the frontal association cortex due to heterozygous 22q11 gene deletion—the genetic foundation of elevated risk for schizophrenia and related psychiatric disorders in 22q11.2 Deletion Syndrome (22q11DS)—reflects cell state-dependent, temporally restricted vulnerability of transcriptionally diverse subsets of intermediate progenitors and neuroblasts. The molecular and cell biological consequences of these evanescent state-dependent changes include divergent proliferative capacity of the most highly proliferative progenitors, enhanced neurogenic gene expression, altered DNA methylation, and increased numbers of immediate neuroblast progeny at peak neurogenesis in fetal frontal cortex of the *LgDel* 22q11DS mouse model. These altered cell states prefigure a post-natal cohort of upper layer PNs generated at the peak of Layer 2/3 PN genesis—not before or after—whose frequencies are significantly diminished and molecular identities are substantially divergent in medial frontal association, but not primary somatosensory or visual cortices. Thus, frontal association cortices in 22q11DS and schizophrenia more broadly may be pathogenic targets due to vulnerabilities of expanded, highly proliferative, transcriptionally dynamic populations of intermediate progenitors that increase L2/3 PN frequency to enhance cortico-cortical connectivity.

## INTRODUCTION

Projection neurons (PNs) in Layers (L) 2/3 of the cerebral cortex are a target of pathology in multiple psychiatric disorders^1–3^. Loss-or gain-of-function mutations in single genes associated with risk for these disorders can modify cortical neurogenesis, including that of L2/3 PNs^4–9^. It remains unclear, however, whether polygenic disease-associated mutations selectively target L2/3-biased progenitors, particularly in association regions where L2/3 PN numbers are amplified to accommodate increased cortico-cortical connectivity^10, 11^. Such progenitor-and region-selective disruption of neurogenesis could compromise subsequent development of L2/3 PN cortico-cortical circuits^12, 13^ whose dysfunction in schizophrenia and other neurodevelopmental disorders is well-established. To resolve this key issue, we analyzed cortical progenitor dynamics and their consequences in medial frontal association as well as somatosensory and visual cortices of the *LgDel* mouse model^14, 15^ of 22q11 Deletion Syndrome (22q11DS), a polygenic developmental disorder accompanied by substantial vulnerability for multiple psychiatric diseases^16–20^.

Most analyses of cortical PN genesis focus on two precursor types: apical progenitors (aPs): slowly dividing ventricular zone (VZ) stem cells, and basal progenitors (bPs): aP-derived transit amplifying precursors in the adjacent subventricular zone (SVZ), that generate primarily L2/3 PNs^10, 11, 21^. Previous work in *LgDel* mice identified reduced bP proliferation in frontal association areas as an antecedent of diminished L2/3 PN frequency, altered connectivity and behavioral deficits due to 22q11 deletion ^22, 23^. Nevertheless, the relationship of these changes to identities, lineage, or states of bPs, their aP precursors, or immediate neuroblast (NB) progeny remains unknown. Diminished 22q11 gene dosage may influence neurogenic capacity of distinct aP or bP subsets thus limiting pathogenic changes to subsets of upper layer progeny, particularly in association cortices where L 2/3 PN numbers are greatest^5, 24^. aP or bP proliferation may be uniformly slowed, yielding consistently fewer L 2/3 PN progeny throughout the period of association cortical upper layer neurogenesis. Alternately, 22q11 deletion may result in divergent temporal, cell-type or region-specific changes in neurogenic capacities that yield diverse changes of L2/3 PN subsets and the circuits they form in a variety of cortical areas.

We found that 22q11 deletion transiently diminishes numbers and alters identities of frontal association—but not primary somatosensory or visual—cortical L 2/3 PNs generated at the peak of upper layer neurogenesis. Divergent cell states of transcriptionally distinct precursor subsets and their NB progeny prefigure these changes. Thus, there is a discrete period of association cortical developmental vulnerability: when intermediate progenitors maximally amplify L 2/3 PN numbers, in an established model of cortical circuit pathology associated with schizophrenia and other psychiatric disorders^20^. Understanding how 22q11-deleted L2/3 PN progenitors are targeted by and recover from this transient disruption may provide novel opportunities to prevent or adjust aberrant cortical circuit differentiation and function that contributes to behavioral pathology in 22q11DS and psychiatric disorders with which it is associated.

## METHODS

### Animals

All animal procedures were approved by Virginia Tech IACUC (#25-249). Male C57Bl/6N mice hemizygous for mmChr. 16 deletion from *Idd* to *Hira*^14^ (*LgDel*) and/or a *Tbr2*^eGFP^ BAC transgene^25^ were crossed with C57Bl/6N WT dams (Charles River). Pregnant dams (morning of plug detection = embryonic day (E) day 0.5) were sacrificed by rapid cervical dislocation on E12.5, 13.5, 14.5, 15.5 or 16.5 and fetuses collected for analysis. For postnatal day (P) 5 animals, delivery was recorded as P0 after daily mid-morning inspection. Tail or limb samples were collected for PCR genotyping (**Supp. Table 1**).

### Histological Preparation

E13.5, 14.5, 15.5 and 16.5 fetuses and P5 pups were fixed with paraformaldehyde and prepared for immunolabeling^23^ with primary antibodies (1°) followed by species-appropriate fluorescent (2°) antibodies (**Supp. Table 2**).

### S-phase labeling

For cell cycle re-entry, EdU (10 mg/kg) was injected intraperitoneally in E12.5, 14.5 and 15.5 pregnant dams and fetuses collected 24 hours later for EdU Click-&-Go^+TM^ labeling. For P5 birthdating, BrdU (50mg/kg)/IdU (70mg/kg) was injected on E12.5 (BrdU)/ 16.5 (IdU), E13.5 (BrdU) /15.5 (IdU), and E14.5 (IdU)/ 16.5 (BrdU). Sections were immunolabeled as described previously ^26–28^ (**Supp. Table 2**).

### Imaging & Analysis

A 100µm wide counting frame from ventricle to pia was overlayed on tiled confocal images (Leica Stellaris 8) of medial frontal association (mFAC), somatosensory (S1) or visual (V1) cortex in coronal sections standardized to the same rostral-caudal locations. Images were counted blind to genotype with counts recorded in overlays for each channel/probe.

### scRNAseq

A single cell library (Parse Biosciences) was generated from dissected E14.5 cortical hemispheres of individual WT (n=5) and *LgDel* fetuses (n=3)/2 litters, or eGFP^+^ cells sorted (Sony SH800 LE-SH800ZFP) from E14.5 *Tbr2*^eGFP/+^ (n=5) and *Tbr2*^eGFP/*LgDel*^ fetuses (n=7)/4 litters. Single cell (sc)RNAseq reads were processed via “split-pipe” (Parse) to collapse reads into uniquely barcoded cells (UBC) expressing genes identified by unique molecular identifiers after normalization via Harmony (UMIs; Seurat with default QC settings). Dimensionality was reduced to 2,000 highly variable genes x 30 principal components, with ribosomal, mitochondrial, and cell cycle genes excluded for clustering. UMAP plots were generated with Leiden clustering at a resolution of 0.8 with minimum distance of 0.3. Quality control, clustering, and visualization was performed with Trailmaker (Parse). Normalized gene matrices for clusters were exported for additional analyses in R (Seurat), Python (scanpy) or Excel. The processed data and novel scripts (for Palantir^29^ and cell cycle^30^) can be accessed at (DOI: 10.5281/zenodo.20707327).

### Bulk RNAseq

5 WT and 5 *LgDel* biological replicates of RNA from sorted *Tbr2*^eGFP+^ cells from 25 E14.5 WT and 26 *LgDel* fetuses/14 litters, RNA integrity (RIN) scores of 10 (Agilent RNA Tapestation) was sequenced to a mean depth of 118.44 ± 59.46 million reads/sample, 150bp paired-end read length. FASTQC quality check, quality trimming and filtering was completed using BBDuk/BBTools. Genome index was built with STAR in linux command-line using Gencode GRCm38/mm10^31^. Reads were mapped with STAR (default parameters, mean unique alignment score: 87.42%±3.78%). Differential expression (DE) analysis was performed using edgeR 4.2.0^32^. Genes with < 15 reads in ≥ 3 samples were removed, and normalization/batch effects accounted for using calcNormFactors. DE genes were identified by FDR of < 0.1. mRNA abundance (TPM) was quantified with DGEobj.utils package using tpm.direct function in R^33^. Gene set enrichment analysis was performed using logFC values as a ranking metric for the fgsea package in R^34^.

### Real-Time PCR

cDNA was synthesized ^35^ from FACS-sorted *Tbr2*^eGFP+^ cell RNA. qPCR (EvaGreen; BioRad) for multiple transcripts (**Supp. Table 1**) was performed on a CFX384 thermal cycler.

### RNAscope

RNAscope probes (Akoya Biosciences) and negative controls for mRNA targets (**Supp. Table 3**) were run with RNA-Protein co-detection ancillary kit (ACD/ Biotechne) on sections co-labeled for Tbr2 (**Suppl. Table 2**).

### Bisulfite sequencing

Genomic DNA was isolated from WT and *LgDel* FACS E14.5 *Tbr2*^eGFP+^ cortical cells (n=6 WT; 10 *LgDel/*8 litters), and samples commercially prepared for bisulfite sequencing (methylcheck^TM^; Zymo Research). Following quality control post-bisulfite primers were designed to cover regions of interest (**Supp. Table 4**), and validated via qPCR using bisulfite-converted control DNA. Genomic DNA samples were bisulfite converted, and target enrichment performed with validated primers on a 48x48 Fluidigm chip. Amplicons were fitted with barcoded adapters to generate a library for next-generation sequencing (Illumina MiSeq). Reads were identified (llumina base-calling software) and analyzed using a proprietary pipeline (Zymo Research). Sequence data was aligned to the reference genome using Bismark software optimized for bisulfite sequencing and methylation calling. Raw sequence data for sequencing/methylation calls can be accessed at (DOI: 10.5281/zenodo.20707327).

### Statistical Analysis

Genotype and age differences in cell frequency, RNAscope counts, and DNA methylation were assessed by two-tailed unpaired t-tests, Chi-Square or Fisher’s exact tests, two-way ANOVA with Holm-Sidak correction for multiple comparisons, or Mann-Whitney U for non-parametric data. Individual data points are plotted on all graphs. Values are presented as means; error bars represent s.e.m; threshold significance is P < 0.05 unless otherwise noted.

### Data availability

Raw sequencing data have been deposited in the NCBI BioProject database under accession number PRJNA1478287. Analysis scripts, worksheets, and supporting files are publicly available through Zenodo (DOI: 10.5281/zenodo.20707327).

## RESULTS

### Selective disruption of bP proliferation and NB generation during peak L2/3 PN genesis

Multiple studies in 22q11-deleted mouse models report L 2/3 PN loss in frontal cortex^22, 23^, and these studies, as well as recent analyses of human cortical organoids, suggest 22q11 deletion disrupts progenitor proliferation^36–38^. 22q11 deletion may render L2/3 PN progenitors proliferatively deficient throughout upper layer neurogenesis, or it may preferentially target temporally distinct progenitor subsets and progeny, thereby disrupting genesis of a limited L 2/3 PN cohort. Thus, we analyzed, aP, bP, NB and <u>e</u>arly differentiating (e)PN identity and proliferative/neurogenic capacity at onset, peak, and decline of bP-mediated upper layer neurogenesis^39, 40^ *in vivo*. We focused on medial frontal association cortex (mFAC) based upon previous observations of L2/3 PN loss in this region^22, 23^. Initial bP genesis from aPs does not appear altered in *LgDel* mFAC at onset of L2/3 PN genesis (E12.5/13.5). WT and *LgDel* aPs re-enter the cell cycle at similar frequencies and VZ/SVZ positions as bPs emerge (E12.5/13.5; **Figure 1a** & **Supp. Fig. 1**). In parallel, E13.5 newly generated Pax6^+^/Tbr2^+^ bP frequencies and distributions are similar in both genotypes (**Figure 1b** & **Supp. Fig. 1**). Apparently, 22q11 deletion does not disrupt initial aP-to-bP yield at the outset of L2/3 PN genesis.

**Figure 1:**
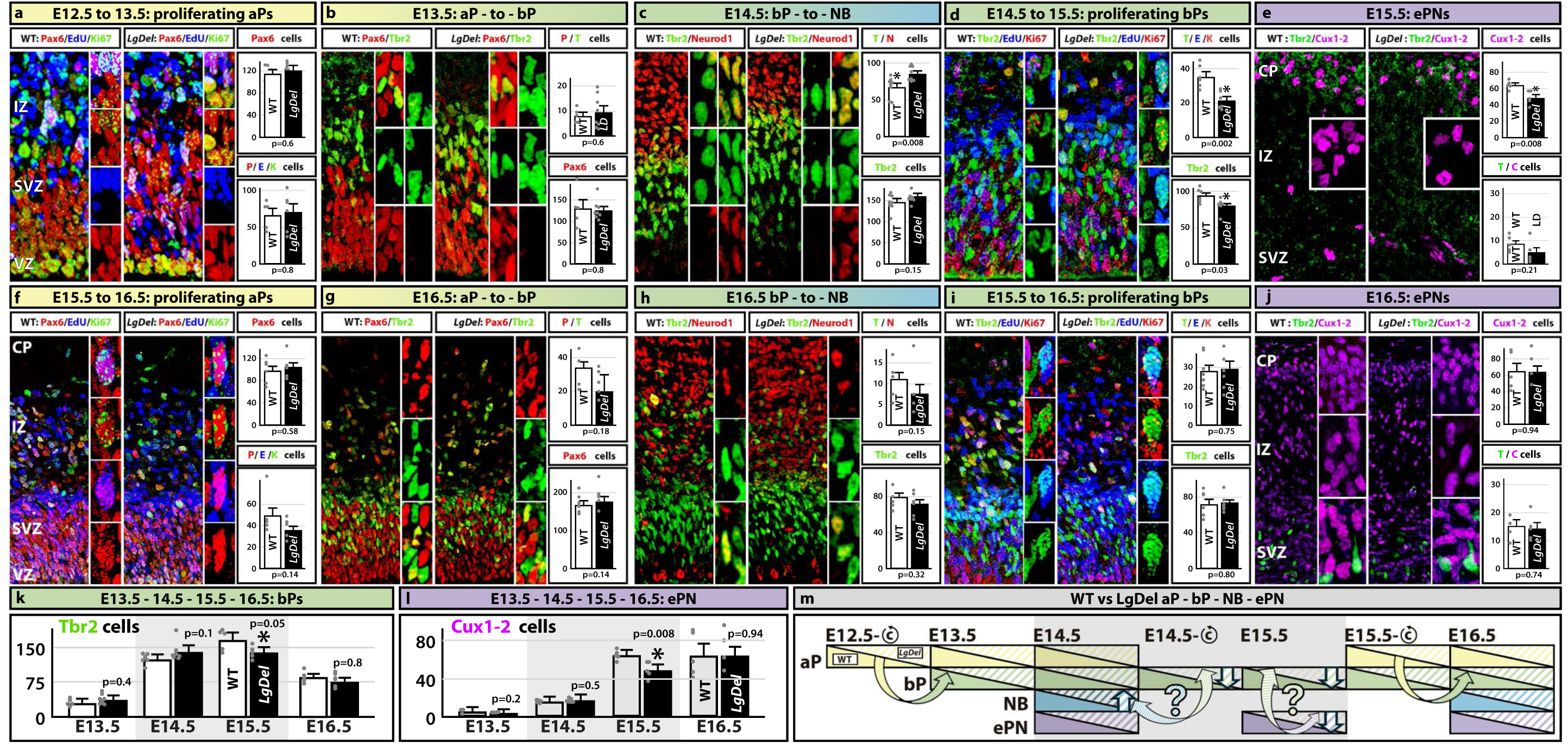
Progenitor dynamics during upper layer neurogenesis in LgDel vs. WT mFAC. **a**. Equivalent frequency of Pax6^+^/EdU^+^/Ki67^+^ aPs re-entering the cell cycle between E12.5 and E13.5 in WT and *LgDel* (unpaired t-test, WT: n = 5, *LgDel*: n = 6). **b**. Equivalent Pax6^+^/Tbr2^+^ newly generated E13.5 bP frequencies in WT and *LgDel* (unpaired t-test WT: n = 5, *LgDel*: n = 8). **c**. Significantly increased *LgDel* Tbr2^+^/Neurod1^+^ presumed E14.5 newly generated NB frequency (unpaired t-test, WT: n = 8, *LgDel*: n = 8). **d**. Diminished cell cycle re-entry of E14.5 *LgDel* Tbr2^+^ presumed bPs (unpaired t-test, WT: n = 6, *LgDel*: n = 6). **e.** Frequency of *LgDel* presumed early differentiating Cux1-2^+^ ePNs in the CP declines significantly at E15.5, (unpaired t-test WT: n = 5, *LgDel*: n = 5). **f.** Frequencies of E15.5 *LgDel* and WT aPs re-entering the cell cycle by E16.5 are equivalent (unpaired t-test WT: n = 5, *LgDel*: n = 5). **g.** Frequency of E16.5 *LgDel* newly generated bPs is equivalent to WT (unpaired t-test WT: n = 7, *LgDel*: n = 7). **h.** Equivalent frequencies of E16.5 newly generated NBs in *LgDel* and WT (unpaired t-test, WT: n = 7, *LgDel*: n = 7). **i.** Equivalent frequencies of *LgDel* E15.5 to 16.5 bPs re-entering the cell cycle (unpaired t-test WT: n = 7, *LgDel*: n = 6). **j.** Equivalent WT and *LgDel* frequencies of E16.5 early differentiating L 2/3 PNs in the CP, and SVZ/IZ Tbr2^+^/Cux1-2^+^ NBs (unpaired t-test, WT: n = 5, *LgDel*: n = 6). **k.** bP frequencies before, during and after peak L 2/3 PN genesis (unpaired t-test E13.5, WT: n = 5, *LgDel*: n = 8; E14.5, WT: n = 6, *LgDel*: n = 7; E15.5, WT: n = 5, *LgDel*: n = 5; E16.5, WT: n = 6, *LgDel*: n = 5). **l.** ePN frequencies throughout, L2/3 PN genesis (E13.5, WT: n = 3, *LgDel*: n = 4; E14.5, WT: n = 6, *LgDel*: n = 7; E15.5, WT: n = 5, *LgDel*: n = 5; E16.5, WT: n = 5, *LgDel*: n = 6). **m.** Summary of aP/bP/NB/ePN dynamics before, during, and after peak L 2/3 PN genesis. *LgDel* and WT aP frequencies and proliferative dynamics appear comparable; however, *LgDel* bP, NB and ePN frequencies and proliferative dynamics diverge between E14.5 and E15.5 (arrows; question marks indicate potential unstable relationships), the peak of L 2/3 PN genesis (gray shading), and then return to WT levels between E15.5 and E16.5.

Proliferative and neurogenic characteristics of the rapidly expanding E14.5 bP cohort diverge substantially in *LgDel* mFAC. The yield of presumed newly generated E14.5 Tbr2^+^/Neurod1^+^ NBs^41, 42^ increases significantly in *LgDel* (20% > WT), without change of Tbr2^+^ cell frequency (**Figure 1c**). In parallel, frequency of E14.5 *LgDel* bPs re-entering the cell cycle by 15.5 declines (40% < WT), accompanied by decreased E15.5 Tbr2^+^ cell frequency (9% < WT; **Figure 1d**). bP and NB locations in the SVZ and intermediate zones (IZ) are preserved in E14.5 *LgDel* mFAC; however, distribution of NBs and cycling bPs change within each zone (**Supp. Fig. 1**). Paradoxially, frequency of cortical plate (CP) E15.5 Cux1-2^+^ ePNs, presumed to be recently generated L2/3 PNs that have completed migration based upon minimal SVZ dual labeling with Tbr2, is significantly diminished in the *LgDel* rather than enhanced as expected due to increased E14.5 NB frequency (**Figure 1e** & **Supp. Fig. 1**).

To determine whether these changes continue as L 2/3 PN genesis declines, we assessed aP and bP dynamics as well as aP, bP NB and ePN frequency between E15.5 and E16.5. The frequency and VZ/SVZ position of *LgDel* aPs re-entering the cell cycle between E15.5 and 16.5 as well as that of newly generated E16.5 bPs are statistically indistinguishable from WT (**Figure 1f, g**). In parallel, frequencies of E15.5 bPs re-entering the cell cycle and E16.5 *LgDel* newly generated NBs are similar in the two genotypes (**Figure 1h,i**). Finally, E16.5 *LgDel* ePN frequency and CP distribution returns to WT levels (**Figure 1j**; **Supp. Fig. 1**). Thus, a transient change in proliferative dynamics in 22q11-deleted mFAC VZ/SVZ selectively depletes bPs, increases NB progeny, and paradoxically diminishes ePN frequency at peak L 2/3 PN genesis, between E14.5 and E15.5 (**Figure 1k,l, m**). As L 2/3 PN genesis declines by E16.5, these 22q11-deletion associated changes in proliferative dynamics and neurogenesis are no longer detected.

### 22q11 deletion disrupts aP, bP and NB cell states during peak L2/3 PN genesis

Our analysis of *in vivo* progenitor/progeny dynamics before, during and after peak L 2/3 PN genesis indicates that cortical progenitors and progeny may be uniquely sensitive to 22q11 deletion during maximal expansion of the L 2/3 PN progenitor cohort between E14.5 and E15.5. It is not clear, however, whether 22q11 deletion-dependent changes target distinct subsets of proliferating bPs, the aPs that generate them, or the NB progeny they yield. Thus, we compared transcriptional identities of bPs, their aP precursors, and NB progeny from *LgDel* and WT E14.5 cortex—when bP genesis of L2/3 PNs reaches peak frequency in WT mice^40^. We analyzed 4772 WT and 3272 *LgDel* cells (median read depth: 120,000 reads/cell): ∼75% from 5 WT and 3 *LgDel* E14.5 cortices; ∼25% from 5 WT and 7 *LgDel* E14.5 *Tbr2*^eGFP+^cell-sorted samples enriched for bPs and immediate progeny^25^. An unbiased UMAP analysis of WT and *LgDel* unsorted and *Tbr2*^eGFP+^ cells identifies discrete interneuron, endothelial, glia-tanycyte, and neuroepithelial clusters flanking aP, bP and NB clusters (**Figure 2a**): two *Pax6/ Nes*^+^/*Fabp7*^+^ /*Slc1a3*^+^ aP subsets, a separate aggregate of three *Tbr2*^+^/*Tis21*^+^/*Neurog2*^+^ bPs, and six contiguous *Neurod1^+^/Dcx^+^/ Tubb3*^+^ NB subsets (**Figure 2a** *<u>inset</u>*; **Supp. Fig. 2**). No aP, bP, or NB subsets are lost or added in *LgDel* (**Figure 2a***, lower panels*). Equivalent aP, bP or NB diagnostic gene expression across WT and *LgDel* subsets reinforces genotypic congruity (**Supp. Fig. 2**). Finally, 21/28 *LgDel*-deleted 22q11 orthologues are expressed at varying levels across all aP, bP and NB subsets and, as expected, levels are diminished by approximately 50% in *LgDel* (**Supp. Fig. 3**).

**Figure 2:**
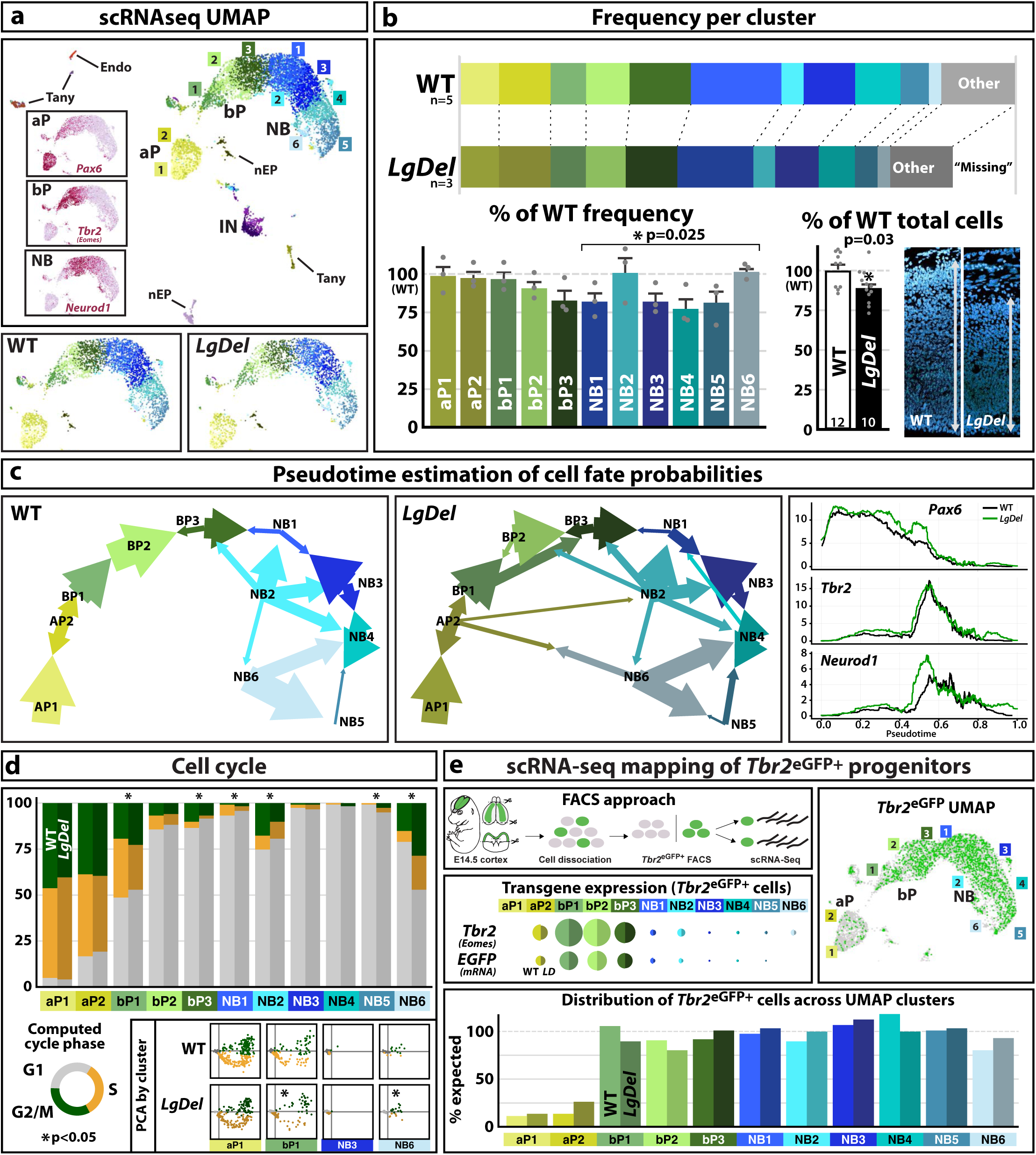
Single cell transcriptome analysis defines divergent cell states among 22q11-deleted cortical progenitors at peak L 2/3 PN genesis. **a**. ***<u>top</u>***: UMAP of single cell RNA sequences (scRNAseq) of 8044 WT and *LgDel* E14.5 cortical cells. Endothelial cells (Endo), tanycytes (Tany), neuroepithelial precursors (nEP) and GABAergic interneurons (IN) cluster discretely. There is an aP aggregate with two subsets (yellow; 1,2), and a continuous bP/NB domain comprising 9 overlapping subsets: 3 bPs (green; 1-3) and 6 NBs (blue; 1-6). ***<u>inset</u>***: *Pax6^+^*, *Tbr2^+^*, and *Neurod1^+^* cells map to aP, bP or early NB domains. ***<u>bottom</u>***: All aP, bP and NB subsets are shared in WT (non-sorted: 3402 cells) and *LgDel* (non-sorted: 1993 cells) E14.5 cortex. **b**. ***<u>top</u>***: E14.5 WT and *LgDel* aPs, bPs, and NBs, normalized to WT levels, vary proportionately (<u>Other</u> = cells outside aP/bP/NB aggregates). ***<u>bottom left</u>***: *LgDel* bP2 and 3 proportions decline, and *LgDel* NB subsets diverge (per animal, Mann-Whitney; n = 5 WT, 3 *LgDel*). ***<u>bottom right</u>***: *LgDel* vs. WT normalized (%WT) *in vivo* cortical cell frequency (unpaired t-test WT: n = 10, *LgDel*: n = 12; arrow-apical to basal; **bottom** left). **c**. ***<u>left & middle</u>***: Pseudo-time analyses of WT vs. *LgDel* transcript frequency and inferred cell fate probabilities. ***<u>right</u>***: Transcript frequency as a function of pseudotime for *Pax6^+^*, *Tbr2^+^*and *Neurod1^+^* cells. **d**. ***<u>top</u>***: G2/M (green), S (orange), or G1 (gray) proportions across aP, bP or NB subsets based upon canonical cell cycle gene expression. G2/M and S cells decline in *LgDel* bP1 and 3 as well as NB 1 and 2, and increase in *LgDel* NB5 and 6 (per animal, Mann-Whitney; n=5 WT, 3 *LgDel*). ***<u>bottom</u>***: PCA of G1 (gray), G2/M (green) and S (orange) cells in WT vs. *LgDel* aP and NB subsets with no differences (aP1, NB3) vs. significant differences (*bP1, *NB6). **e**. ***<u>top left</u>***: Fluorescence-activated cell sorting of *Tbr2*^eGFP+^ cells to enrich bPs and immediate NB progeny. ***<u>middle left</u>***: Similar frequencies of *Tbr2* and *eGFP* transcripts across aP, bP and NB subsets in WT and *LgDel*. ***<u>top right</u>***: WT and *LgDel Tbr2*^eGFP+^ cells (green dots) across UMAP. ***<u>bottom</u>***: similar frequencies of *Tbr2*^eGFP+^ cells in WT and *LgDel* aP, bP and NB subsets.

Cell frequencies (non-sorted only), normalized to WT, in the two aP and first two bP subsets are proportionally equivalent in *LgDel* and WT. Diminished cell frequency in the 3^rd^ *LgDel* bP subset, relative to WT, does not reach statistical significance (**Figure 2b**). There is, however, a significant decline across *LgDel* bP 3 as well as NB1, 3, 4 and 5 subsets to approximately 75% of WT. This decline predicts diminished cortical cell density and thickness due to “missing” cells, a change consistent with *in vivo* E14.5 *LgDel* vs. WT mFAC morphology (**Figure 2b**, *inset*). Inferred trajectories of *LgDel* vs. WT aP, bP and NB subsets, based upon pseudotime analysis^29^, also differ substantially (**Figure 2c**), suggesting instability of cell states among progenitors and progeny during peak L2/3 PN genesis. In WT, aP1 progresses to aP2 and then serially to 3 bP subsets that precede somewhat more labile NB progeny, some of which retain reciprocal relationships to bP3 as well as one another (**Figure 2c**, *left*). In contrast, *LgDel* aPs contribute to all bP and NB subsets, the serial trajectory of bPs is altered, and NBs have reciprocal relationships with multiple aP and bP subsets (**Figure 2c**, *left*). *Pax6* relative expression increases across pseudotime, coincident with divergence *Tbr2* and *Neurod1* expression in register with apparent *LgDel* aP/bP/NB lability (**Figure 2c**, *right*). In parallel, several *LgDel* bP and NB subsets have variable proportions of cycling cells (**Figure 2d**) based upon frequency of G1-, S-, and G2/M-associated transcripts^30^. Inferred proliferative frequency declines in *LgDel* bP 1 & 3 and NB1 &2 subsets, suggesting diminished self-renewal or enhanced neurogenesis. In contrast, NB5 & 6 proliferative frequencies increase in parallel with altered aP2 relationships (**Figure 2c**). Thus, despite populating similar subsets as WT, E14.5 *LgDel* bP/NB cell frequencies, inferred lineal relationships, and proliferative states diverge from WT counterparts.

Divergent *LgDel* E14.5 bPs and their immediate NB progeny may be distributed uniformly across all aP, bP and NB populations shared by both genotypes, or restricted to distinct subsets. To determine whether recently-generated bPs and NBs segregate in transcriptionally defined subsets, we sorted presumed *LgDel* and WT E14.5 bPs and their immediate progeny labeled by a *Tbr2*^eGFP^ reporter^25^ and mapped them onto the scRNAseq data (**Figure 2e**). *LgDel* and WT *Tbr2*^eGFP+^ bPs and NBs, which express *eGFP* and *Tbr2* transcripts at equivalent levels, map onto bP1 - 3 and NB1 - 6 at approximately equivalent frequencies, with little aP presence (**Figure 2e**, *bottom*). Thus, neither *LgDel* nor WT E14.5 bPs and related recently generated NBs or ePNs segregate within defined subtypes. Instead, they distribute broadly across multiple transcriptional and proliferative states.

### Transcriptional divergence in 22q11-deleted bPs and their NB progeny

Molecular differences among related cohorts of bPs, NBs and newly generated neurons at peak bP-mediated L2/3 PN genesis may not be fully resolved by scRNAseq. Thus, we performed bulk RNA sequencing on FAC-sorted *LgDel* and WT E14.5 *Tbr2*^eGFP+^ cortical cells that constitute a lineally related sub-population across multiple bP and NB subsets (see **Figure 2**). Based upon 5 pooled *Tbr2*^eGFP+^sorted samples/genotype (E14.5 WT: 25, *LgDel*: 26 *Tbr2*^eGFP^cortices/14 litters), 19645 genes are significantly expressed (≥ 15 read counts) in WT, and 18057 in *LgDel Tbr2*^eGFP+^ cells (**Figure 3a, b**). Of these, 79 are differentially expressed (DE, FDR < 0.1). Fifty DE genes are upregulated, thirteen of which are associated with accelerated neurogenesis^43–49^ (**Figure 3b**, *inset*); twenty-nine are downregulated, including twenty-four 22q11 genes (**Figure 3a, b**). 8/50 Hallmark gene sets^50^ associated with cell cycle, metabolic and cytoskeletal pathways were significantly enriched (p. adj.<0.05) among DE genes (**Supp. Fig. 4**). In parallel, many DE genes— including the 22q11-deleted subset—are selectively expressed in the E14.5 VZ, SVZ, or IZ; many are enhanced in the SVZ ^30^, and some differ substantially in expression level across the anterior-posterior cortical axis (**Supp. Fig. 5**). Quantitative (q)PCR of a subset of 22q11 and DE genes in *Tbr2*^eGFP+^ FACS samples provides initial validation (**Supp. Fig. 5**) as does detection at varying levels across scRNAseq bP and NB subsets where *Tbr2*^eGFP+^ cells are found (**Supp. Fig. 6**).

**Figure 3:**
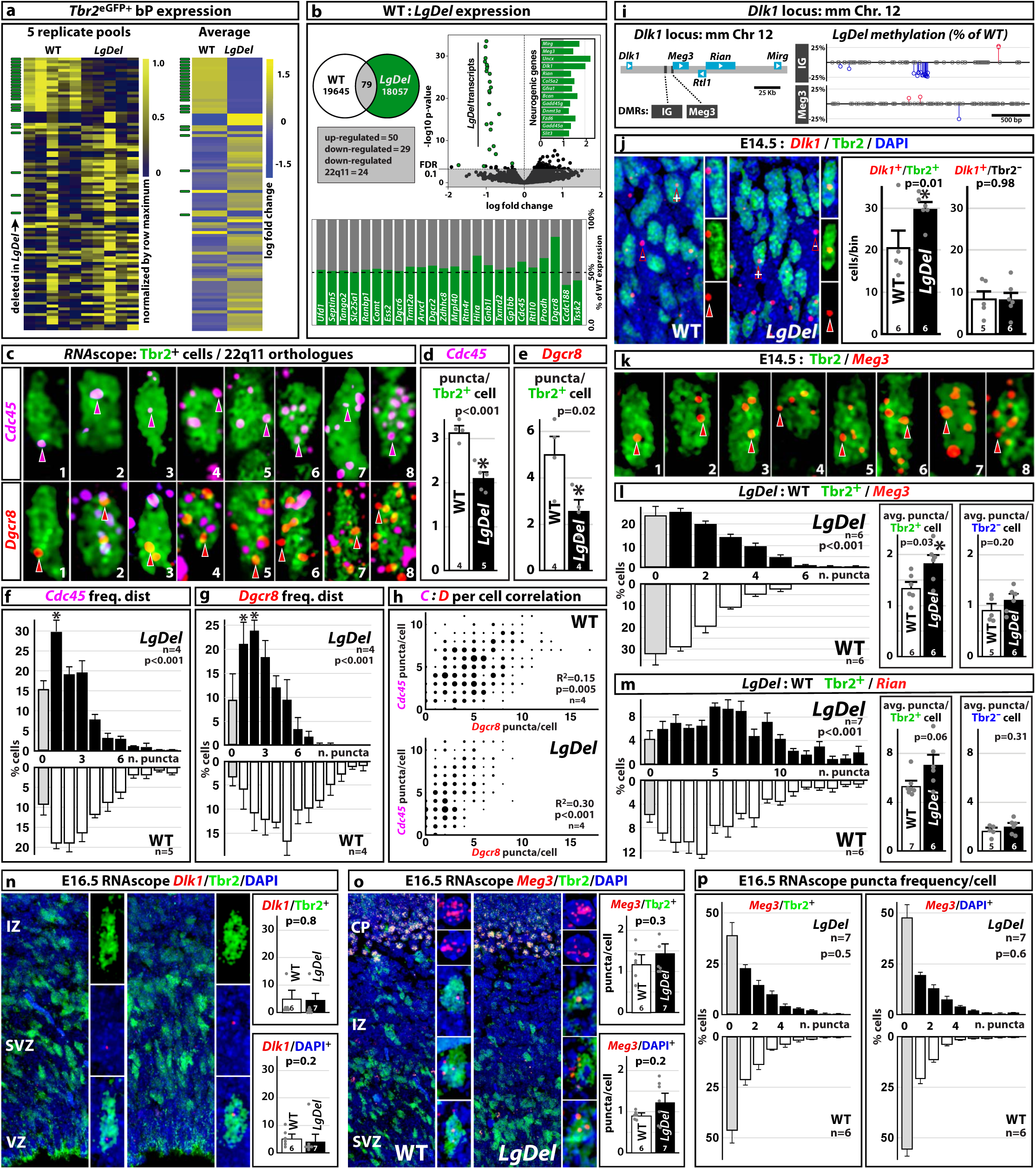
Differential gene expression in 22q11-deleted E14.5 bPs and bP progeny. **a**. Individual (***left***) and summary (***right***) heat maps of differentially expressed (DE) genes in 5 pooled samples of *Tbr2*^eGFP+^ cells (green bars, *far left* = 22q11-deleted genes). **b**. DE gene volcano plot, including 13 up-regulated DE genes associated with cortical neurogenesis (*<u>inset</u>*) and 24 down-regulated 22q11-deleted genes (green dots, left). **c**. RNAscope puncta for 22q11-deleted genes *Cdc45* (violet, arrowheads) and *Dgcr8* (red, arrowheads) in Tbr2^+^ cells (green)in E14.5 WT and *LgDel* VZ/SVZ. Numbers (lower right) indicate puncta counted per each illustrated cell. **d**. *Cdc45* puncta frequency declines significantly in *LgDel* Tbr2^+^ cells (unpaired t-test WT: n = 4, *LgDel*: n = 5). **e**. *Dgcr8* puncta frequency declines significantly in *LgDel* Tbr2^+^ cells (unpaired t-test WT: n = 4, *LgDel*: n = 4). **f**. The distribution of *Cdc45* puncta in Tbr2^+^ cells in *LgDel* vs. WT includes more Tbr2^+^ cells with no puncta (gray bars, *left*) and fewer Tbr2^+^ cells with the highest puncta frequencies. **g.** The distribution of *Dgcr8* puncta frequencies includes more Tbr2^+^ cells with no *Dgcr8* puncta (gray bars, *left*), and fewer Tbr2^+^cells with greater puncta frequency. **h**. Minimal correlation of *Cdc45* and *Dgcr8* puncta frequencies/Tbr2^+^ cell in WT (*top*) and *LgDel* (*bottom*). **i**. Transcripts (light blue) and differentially methylated regions (DMRs, black boxes) in the murine *Dlk1* locus (Chr. 12; *left*). The IG, but not *Meg3* DMR is significantly demethylated (2-way ANOVA, WT: n = 6, *LgDel*: n = 10). **j**. ***left***: Single *Dlk1* puncta in E14.5 WT and *LgDel* Tbr2^+^ (green “+” arrowheads) and Tbr2^-^(blue, “-“ arrowheads) cells. ***right***: *LgDel Dlk1*^+^/Tbr2^+^ cells (only single *Dlk1* puncta seen) increase significantly (unpaired t-test WT: n = 5, *LgDel*: n = 6). **k**. *Meg3* puncta (red, arrowheads) in Tbr2^+^ cells (green) of WT or *LgDel* E14.5 SVZ. Numbers (*lower right*) indicate puncta count for each illustrated cell. **l.** Distribution and frequencies of *Meg3* puncta in *LgDel* and WT Tbr2^+^ and Tbr2^-^cells. *Meg3* puncta frequency increases in Tbr2^+^ cells but not Tbr2^-^cells (*<u>insets</u>, right*: Tbr2^+^: Unpaired t-test WT: n = 6, *LgDel*: n = 6, Tbr2^-^: Unpaired t-test WT: n = 5, *LgDel*: n = 6). **m**. Distribution and frequencies of *Rian* puncta in *LgDel* and WT Tbr2^+^ and Tbr2^-^cells. *Rian* puncta frequency increases in Tbr2^+^ but not Tbr2^-^cells (*<u>insets</u>, right*: Tbr2^+^: unpaired t-test WT: n = 7, *LgDel*: n = 6, Tbr2^-^: unpaired t-test WT: n = 5, *LgDel*: n = 6). **n**. *Dlk1* in Tbr2^+^ and Tbr2^-^cells in E16.5 WT and *LgDel* VZ/SVZ/IZ is limited to 1 punctum per Tbr2^+^ or Tbr2^-^cell. Compared to E14.5, *Dlk1*^+^/Tbr2^+^ and /Tbr2^-^cell frequencies decline and are statistically indistinguishable in the two genotypes. **o**. *Meg3* puncta frequencies in Tbr2^+^ and Tbr2^-^(DAPI^+^) are statistically indistinguishable in E16.5 WT and *LgDel* SVZ/IZ (CP not quantified). **p**. Equivalent frequency distributions of *Meg3* RNAscope puncta/cell in E16.5 WT vs. *LgDel* Tbr2^+^ (***top***) and Tbr2^-^(***bottom***) SVZ/IZ cells (Levene’s F test).

**Figure 4:**
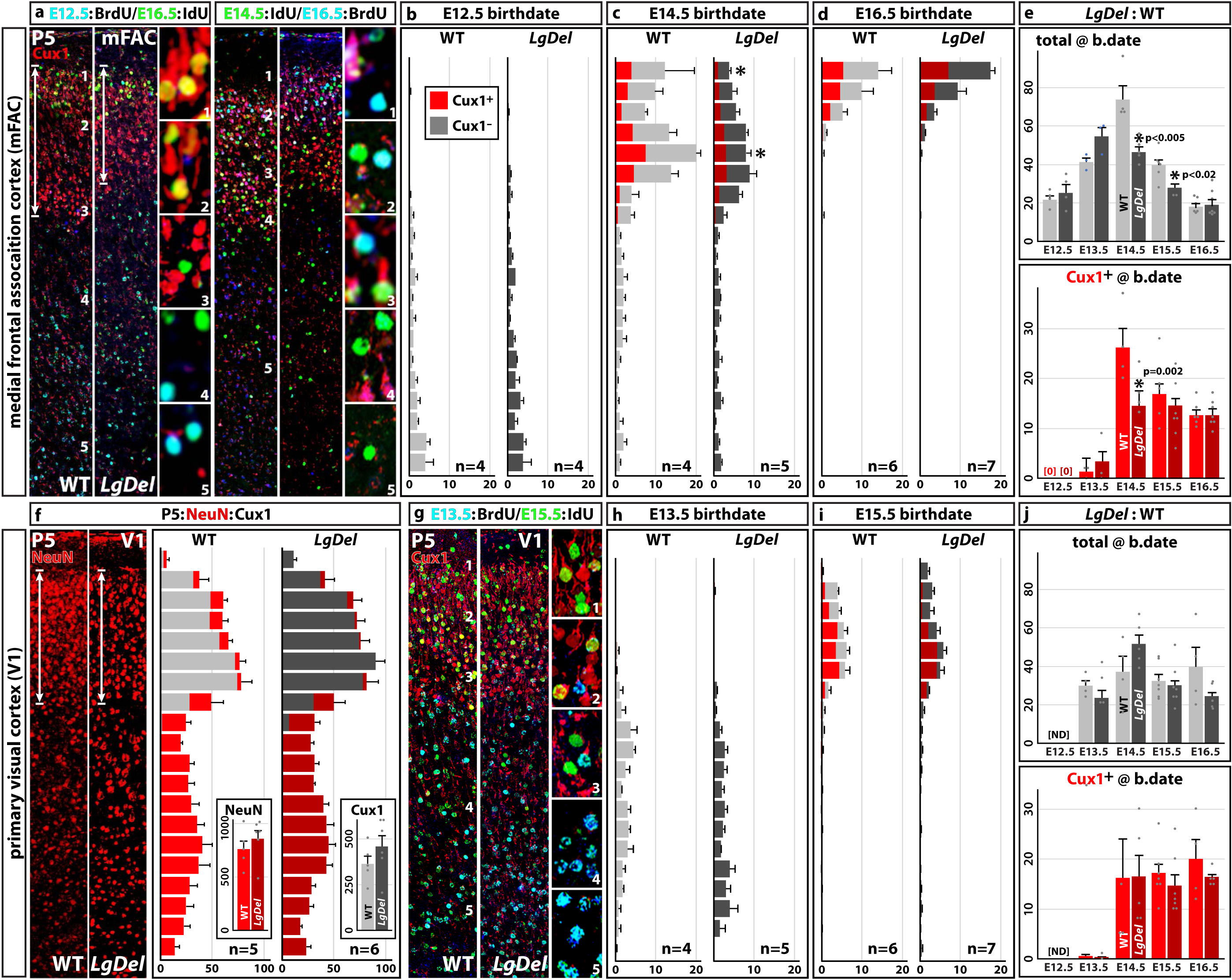
Transient, region-selective diminished L 2/3 PN genesis in postnatal frontal association cortex. **a.** mFAC L2/3 PNs (Cux1^+^) birthdated by dual S-phase markers. Brackets and arrows (***far left***) indicate *LgDel* vs. WT L 2/3 attenuation. P5 E12.5-generated lower layer cells (BrdU^+^, cyan) are distinct from E16.5-generated upper layer cells (IdU^+^, green) including Cux1^+^ L 2/3 PNs (red, ***left***). E14.5-generated (IdU^+^, green) as well as E16.5-generated (BrdU^+^, cyan) upper layer cell include Cux1^+^ L 2/3 PNs (***right***). <u>Panels 1-5</u>: ***<u>left</u>***: E16.5 Cux1^+^/IdU^+^ or IdU^+^ upper layer cells (locations: 1, 2, 3, left) and E12.5 Cux1^-^/BrdU^+^ lower layer cells (locations 4,5). ***<u>right</u>***: E14.5-(green) vs. E16.5-(cyan) generated Cux1^+^ (red) and Cux1^-^cells in upper (1-4) and, more rarely, lower (5), layers at higher magnification. **b**. Similar distribution of *LgDel* and WT E12.5-generated BrdU^+^/Cux1^−^ cells across 20 equivalent bins (ventricle to pia); no E12.5-generated BrdU^+^/Cux1^+^ cells are detected. **c**. Upper layer location of E14.5-generated *LgDel* vs. WT IdU^+^/Cux1^+^ and IdU^+^/Cux1^−^ L2/3 PN cohorts is similar; however, frequencies and distributions diverge. **d**. *LgDel* vs. WT E16.5-generated BrdU^+^/Cux1^+^ and Cux1^−^ cells frequencies and distributions are statistically indistinguishable. **e**. ***top:*** Diminished L2/3 PN genesis is not detected at E12.5 or 13.5 in LgDel; it reaches a significant maximum at E14.5, continues through E15.5 and is no longer detected at E16.5. Selective loss of the Cux1^+^ L 2/3 PN subset (***bottom***) in *LgDel* is restricted to E14.5 (2-way ANOVA with Šídák’s multiple comparison test). **f**. Cortical neuron frequency and distribution, based upon NeuN^+^ cells in all cortical layers, is statistically indistinguishable in WT (*left*) and *LgDel* (*right*) primary visual cortex (V1). **g**. Dual E13.5 BrdU(cyan)/IdU E15.5 (green) labeling in WT and *LgDel* V1. **h**. Statistically indistinguishable frequency/distribution of E13.5 BrdU as well as BrdU/Cux1^+^ cells in WT vs. *LgDel* V1. **i**. Statistically indistinguishable frequency/distribution of E15.5 IdU as well as IdU/Cux1^+^ cells in WT vs. *LgDel* V1. **j.** Neurogenesis in V1 of WT vs. *LgDel* is comparable from E13.5 through E16.5 (*top*) as is genesis of Cux1^+^ presumed L 2/3 PNs (*bottom*).

To confirm the accuracy of these data, we completed a parallel analysis of mRNA frequency in E14.5 *LgDel* and WT Tbr2+ and Tbr2^-^SVZ cells using RNAscope quantitative *in situ* labeling. As initial calibration, we assessed transcript frequency/cell, which is independent of cell size (**Supp. Fig. 7**) for two 22q11 genes with selective SVZ expression: *Cdc45* and *Dgcr8*. As expected, labeling for both transcripts declines by approximately 50% in individual *LgDel* mFAC Tbr2^+^ and Tbr2^-^(DAPI^+^) SVZ/IZ cells (**Figure 3c**-**e**; **Supp. Fig. 7**). These changes, however, do not reflect shifted means of normally distributed *Cdc45* or *Dgcr8* transcripts/cell in *LgDel* vs. WT. Instead, maximal *Cdc45* or *Dgcr8* puncta frequency in each genotype falls between 1 and 3/cell. The 50% decline reflects more *LgDel* Tbr2^+^ and Tbr2^-^cells with no *Cdc45* or *Dgcr8* puncta, and fewer cells with highest puncta counts (**Figure 3f, g**; **Supp. Fig. 7**). These changes are apparently independent, since there is minimal correlation of *Cdc45* vs. *Dgcr8* puncta /cell in Tbr2^+^ or Tbr2^-^SVZ/IZ cells of either genotype (**Figure 3h**; **Supp. Fig. 7**). This spectrum of divergent *LgDel* transcriptional states indicated by varying levels of 22q11 gene expression—from WT to apparently “null”—suggests substantial dynamism among bPs and neighboring cells.

To assess DE gene specificity, we analyzed genomic regulation and quantitative expression of three cortical neurogenesis-associated genes: *Dlk1*, *Meg3* and *Rian* (see **Figure 3b**, *inset*)^51–53^ in Tbr2^+^ and Tbr2^-^VZ/SVZ/IZ cells. These transcripts, plus *Rtl1* and *Mirg*, are encoded by the multigene *Dlk1* locus on mmChr12^54^ (**Figure 3i**). Two *Dlk1* locus-control regions^54^ can be differentially methylated: an intergenic region (IG-DMR) that influences the entire locus and an adjacent region that influences *Meg3* (**Figure 3i**, *left*). We evaluated *Dlk1* locus methylation in 5 biological replicates of E14.5 FACS-sorted Tbr2^eGFP+^ cells. The *LgDel* IG-DMR, but not *Meg3*-DMR, is hypomethylated at multiple CpGs (**Figure 3i**, *right*), suggesting that *Dlk1* locus transcripts may be up-regulated in parallel in *LgDel* Tbr2^+^ VZ/SVZ cells *in vivo*. *Rtl1* and *Mirg* fall beneath detection threshold (FDR <0.1) in both genotypes; however, *Dlk1*, *Meg3* and *Rian* increase by approximately 2-fold in *LgDel Tbr2*^eGFP+^ cells based upon qPCR (**Supp. Fig. 6**). *Dlk1*, *Meg3* and *Rian* transcripts are detected only in bP and NB scRNAseq subsets, further validating the bulk seq data (**Supp. Fig. 6**). *In vivo*, *Dlk1* is limited to 1 punctum per WT and *LgDel* Tbr2^+^ or Tbr2^-^cell (**Figure 3j**, *left*). Thus, the 2-fold *Dlk1* increase in *LgDel* Tbr2^+^ but not Tbr2^-^cells reflects greater *Dlk1*^+^/Tbr2^+^ cell frequency (**Figure 3j**). *Meg3* and *Rian* puncta/cell increase approximately 2-fold per Tbr2^+^, but not Tbr2^-^cell; however, *Rian* is marginally significant (**Figure 3k**-**m**). Increased *Meg3* and *Rian* puncta in Tbr2^+^ cells reflects unequal variance: *Meg3^-^*or *Rian^-^*Tbr2^+^ cell frequency decreases and Tbr2^+^ cell frequencies with the most puncta increase (**Figure 3l, m**). Apparently, dynamic, variable neurogenic gene expression in E14.5 22q11-deleted Tbr2^+^ VZ/SVZ cells *in vivo* parallels altered bP/NB cell states at peak L 2/3 PN genesis.

To determine whether enhanced expression of *Dlk1* locus genes is retained after peak L 2/3 PN genesis, as bP, NB, and ePN frequencies return to WT (see **Figure 1**), we quantified *Dlk1* and *Meg3* mRNA in E16.5 *LgDel* and WT mFAC Tbr2^+^ and Tbr2^-^VZ/SVZ/IZ cells. Tbr2^+^ and Tbr2^-^*Dlk1^+^* cell frequency—still 1 punctum/cell—is equivalent in *LgDel* and WT E16.5 VZ/SVZ (**Figure 3n**), despite substantial decline in *Dlk1^+^* cell frequency from E14.5 levels in both genotypes. E16.5 *LgDel Meg3* puncta/Tbr2^+^ and Tbr2^-^cell frequency distribution are indistinguishable from WT (**Figure 3o, p**). Apparently, when neurogenic dynamics diverge at peak L 2/3 PN genesis in *LgDel* mFAC, there is differential expression as well as cell type-selective genomic regulation of a subset of neurogenic genes that returns to a WT state as L 2/3 PN genesis declines.

### Selective disruption of 22q11-deleted bP states targets genesis of a distinct L2/3 PN cohort

Aberrant precursor dynamics, neurogenic states, a surfeit of E14.5 NBs and a modestly depleted bP pool by E15.5 predicts an inflated peak frequency of E14.5-generated upper layer neurons in post-natal *LgDel* mFAC, followed by substantial decline. Thus, we compared L2/3 PN birthdates in *LgDel* P5 mFAC as postnatal cortical circuit maturation accelerates^55, 56^. There are no differences in E12.5 or E13.5 birthdated, primarily lower layer, cell frequencies in the two genotypes (**Figure 4a**, **b, e**; **Supp. Fig. 8**). In contrast, *LgDel* E14.5 birthdated cell frequency, presumably from the peak proliferative bP cohort that generates an NB surplus (see **Figure 1**), *<u>declines</u>* significantly in *LgDel* (40% < WT; **Figure 4a**, **c**, **e**), a decline that continues as L 2/3 PN genesis slows to half of peak level by E15.5 (**Figure 4e**; **Supp. Fig. 8**). E14.5-and 15.5 generated neurons are found throughout the upper layers; however, in *LgDel*, there is local variation in the distribution within L 2/3 (**Figure 4c**; **Supp. Fig. 8**). As L2/3 PN genesis slows further at E16.5 from E14.5 peak levels, frequency and distribution of *LgDel* E16.5 birthdated cells, exclusively in upper layers, returns to WT (**Figure 4d, e**). Finally, frequency of the E14.5 generated Cux1^+^ subset of *LgDel* L L2/3 PNs^40, 57^ declines significantly (47% < WT; p < 0.002) but not that of E12.5, 13.5, 15.5 or E16.5 (**Figure 4e**, *top*; **Supp. Fig. 8**). Paradoxically, the surfeit of *LgDel* E14.5 NBs prefigures a substantial, transient *<u>decline</u>* in E14.5 birthdated P5 cortical cells including L 2/3 PNs. This neurogenic deficit recovers by E15.5, consistent with apparent restoration of typical proliferative dynamics between E15.5 and E16.5 (see **Figure 1**).

To determine whether transient divergence in L2/3 PN genesis is limited to mFAC or impacts peak L 2/3 PN genesis throughout the cortex, we analyzed L 2/3 PN birthdates in primary somatosensory (S1) and visual cortices (V1). Confirming previous observations^22^, we found no difference in S1 neurogenesis at E14.5 or E16.5 (**Supp. Fig. 9**). To evaluate upper layer neurogenesis across the anterior-posterior cortical axis, we also analyzed neuron frequency, distribution and birthdates in primary visual cortex (V1). Frequencies and distribution of *LgDel* vs. WT NeuN/RbFox3^+^ neurons across all V1 cortical layers is statisitically indistinguishable (**Figure 4f**) as is the case in S1^22^. To assess potential neurogenic divergence despite comparable neuron frequencies, we analyzed birthdates throughout the period of V1 upper layer PN genesis (E13.5, 14.5, 15.5, 16.5; **Figure 4g, h**; **Supp. Fig. 8**). We found no differences in frequency or distribution of *LgDel* vs. WT V1 birthdated cells at these ages (**Figure 4j**, *top*). Similarly, Cux1^+^ L2/3 PN frequency and distribution was indistinguishable for each birthdated cohort in the two genotypes (**Figure 4j**, *bottom*). Apparently, transient disruption of L 2/3 PN genesis during peak upper layer neurogenesis due to heterozygous 22q11 deletion in the *LgDel* mouse is selective for frontal association vs. primary somatosensory or visual cortices.

### Selective divergence of cortical neuron identities due to 22q11 deletion

To resolve whether 22q11-deletion has a broader impact on mFAC or V1 L2/3 PN identities—independent of time of genesis—we analyzed position and frequency of WT and *LgDel* P5 L2/3 PNs identified by canonical makers Cux1-2, Satb2 and Brn2^58–61^ (**Figure 5**). In *LgDel* P5 mFAC, cells labeled by all three markers comprise an attenuated L 2/3 (**Figure 5a**, *brackets***; Figure 5b, c**). mFAC Cux1-2^+^ cells are limited to upper layers; Satb2^+^ cells are enhanced in L 2/3 but found in all layers, and Brn2a^+^ cells are primarily, but not exclusively, in upper layers (**Figure 5a-c**). In *LgDel* mFAC, Cux1-2^+^ PN frequency increases slightly but not significantly (**Figure 5d**, *top*); Satb2^+^ L 2/3 PN frequency increases substantially (Satb2^+^: 194% > WT; **Figure 5d**, *top*), consistent with increased frequency of interhemispheric projecting L 2/3 PNs in *LgDel* frontal cortex^62^; Cux1-2^+^/Satb2^+^ cell frequency declines modestly, but significantly (11% < WT, p=0.05; **Figure 5d**, *middle*); Brn2^+^ upper layer PN frequency is unchanged (**Figure 5d**, *bottom*; **Supp. Fig. 10**), and mFAC lower layer Satb2^+^ PN frequency is equivalent in the two genotypes (**Figure 5b**, *inset*). In contrast, *LgDel* V1 L 2/3 is neither visibly nor quantitatively diminished (**Figure 5e**, *brackets*; **Figure 5f, g**). *LgDel* V1 Cux1-2^+^, Satb2^+^ and Brn2^+^ cell frequency and distribution is comparable to WT (**Figure 5e-h**; **Supp. Fig. 10**). as is Cux1-2^+^/Satb2^+^ L2/3 PN frequency and distribution (**Figure 5f**; **Figure 5h**, *middle*). Thus, broader L 2/3 PN identity changes accompany transient disruption of L 2/3 PN genesis in mFAC; however, these changes are not seen in V1 where 22q11 deletion does not detectably alter genesis or frequency of L 2/3 PNs.

**Figure 5:**
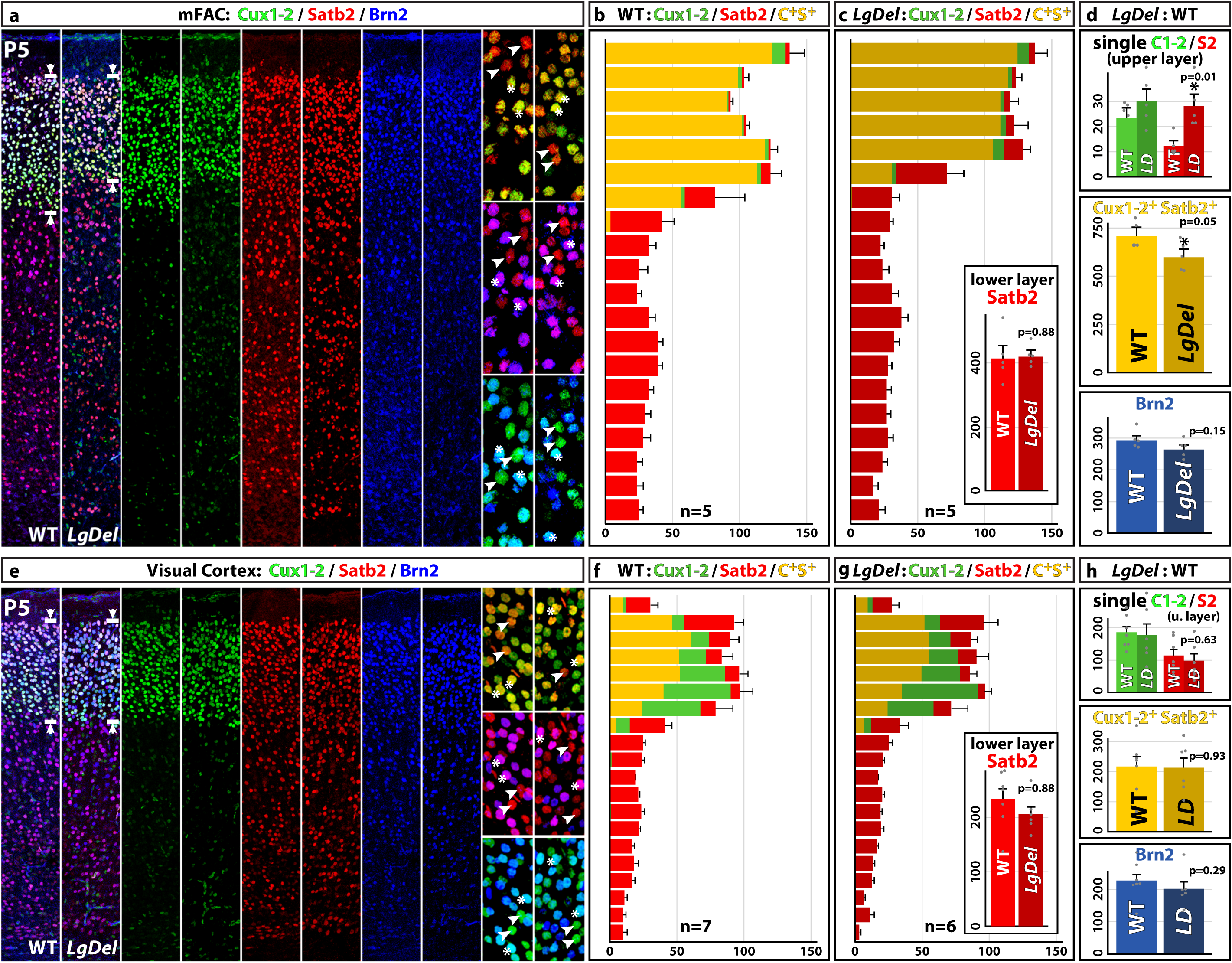
Altered L 2/3 PN identities of in early post-natal 22q11-deleted mFAC and V1. **a**. Cux1-2^+^, Satb2^+^ and Brn2^+^ cells in P5 WT and *LgDel* mFAC; L2/3 attenuation (brackets/arrowheads; **left**). Mosaic distribution of WT and *LgDel* L2/3 single-(arrowheads, ***right***) and double-labeled PNs (asterisks, *right*). **b, c**. Frequency and distribution of single-(green: Cux1-2^+^; red: Satb2^+^) and double-labeled L2/3 PNs (yellow) differs in WT (**b**) vs. *LgDel* (**c**). *<u>inset</u>*: Similar WT and *LgDel* frequencies of lower layer Satb2^+^ cells. **d**. Differences in *LgDel* single (***top***) and double-labeled Cux1-2^+^/Satb2^+^ (***middle***) but not Brn2^+^ single (***bottom***) labeled cells (unpaired t-test). **e**. Cux1-2^+^, Satb2^+^ and Brn2^+^ cells in P5 WT and *LgDel* V1; L2/3 widths are similar (brackets/arrowheads; ***left***). Mosaic distribution of WT and *LgDel* L2/3 single-(arrowheads, ***right***) and double-labeled PNs (asterisks, ***right***). **f, g**. Frequency and distribution of single-(green: Cux1-2^+^; red: Satb2^+^) and double-labeled L2/3 PNs (yellow) differs in WT (**b**) vs. *LgDel* (**c**). *<u>inset</u>*: Similar WT and *LgDel* frequencies of lower layer Satb2^+^ cells. **h**. Differences in *LgDel* single (*top*) and double-labeled Cux1-2^+^/Satb2^+^ (*middle*) but not Brn2^+^ single (*bottom*) labeled cells (unpaired t-test).

## DISCUSSION

The most rapidly dividing, maximally expanding populations of intermediate progenitors biased to generate L2/3 PNs in cortical association areas appear selectively vulnerable to polygenic changes associated with 22q11DS^63, 64^ (**Figure 6**). This vulnerability is not limited to a discrete intermediate progenitor or neuroblast molecular class; instead, it reflects dynamic cell states across molecular marker-defined subtypes^65^. Heterozygous 22q11 gene deletion destabilizes these cell states at peak intermediate progenitor proliferation and L2/3 PN neurogenic division, not before or after (**Figure 6a**). This temporally and regionally selective disruption of frontal association cortical L2/3 PN genesis may contribute to cortical circuit pathology in 22q11DS (**Figure 6b**). Similar disruptions may underlie pathogenesis in schizophrenia and other neurodevelopmental disorders^66, 67^. In typically developing brains, however, dynamic cell states may maximize L 2/3 PN diversity ^68, 69^ by enhancing neurogenic flexibility for rapidly expanding, rapidly dividing progenitors in association cortices of multiple species including humans^70–72^.

**Figure 6:**
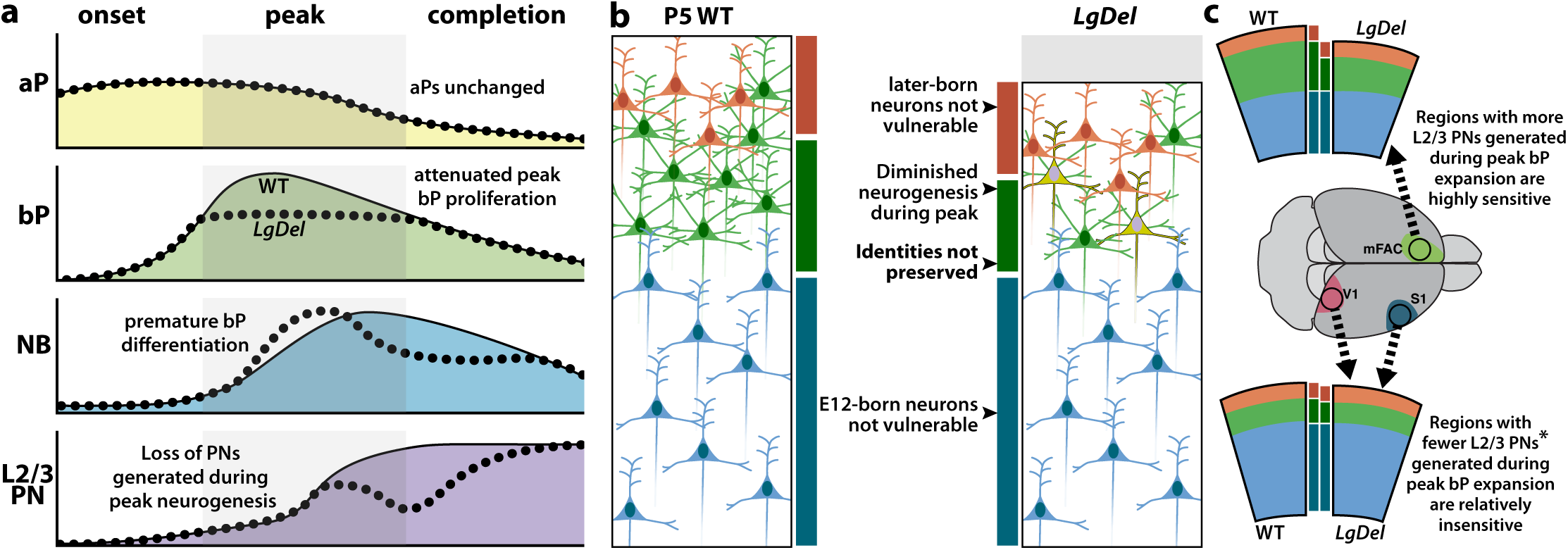
Selective targeting of rapidly expanding, rapidly dividing intermediate progenitor and neuroblast cell states in association cortices due to heterozygous 22q11 deletion. **a**. Schematic of enhanced cell state-dependent vulnerability of rapidly dividing, maximally expanding neurogenic intermediate progenitors in WT (solid lines) and *LgDel* (dotted lines) during onset, peak and completion of association cortex L2/3 PN genesis. Proliferation of aPs is unchanged (***top***); however, intermediate progenitor expansion and variable states among bPs (***upper middle***) and their immediate NB progeny (***lower middle***) during this time define peak vulnerability for L 2/3 PN neurogenic deficits (***bottom***) due to heterozygous 22q11 gene deletion. **b**. Schematic of divergent frequency and identities of 22q11-deleted L 2/3 PNs generated at peak L 2/3 PN neurogenesis—not before or after—in mFAC. Transient disruption of maximal bP-mediated L 2/3 PNs results in a L 2/3 mosaic with as well as a mixture of PNs with divergent identities (gray) as well as altered proportions of birthdates during the final steps of association cortico-cortical circuit development. **c.** Regionally distinct consequences for disrupted peak intermediate progenitor L 2/3 PN genesis in medial frontal association vs. primary sensory and visual cortices in 22q11-deleted mice.

A broad range of polygenic changes, thought to be primary genetic drivers of NDDs^63, 64^, including copy number variants beyond 22q11.2^73, 74^ and combinations of single risk variants^75, 76^ may modulate cell-state dynamics leading to association cortical vulnerability at peak neurogenesis. Such polygenic changes are thought to enhance replication stress in stem cells and related intermediate progenitors^77–79^ to substantially change frequency, viability and identity of rapidly dividing progenitors and their progeny. Replication stress due to 22q11 deletion may reflect diminished dosage of two 22q11 deleted genes that regulate DNA replication: *HIRA* and *CDC45*^24^; both expressed at significant levels in bPs and NBs. Diminished dosage of additional genes in the minimal critical deleted region, including *Ranbp1*^5^, *Dgcr8*^8^, and a subset of mitochondria-localized genes^9^—also expressed in bPs and NBs and associated with neurogenesis in homozygous mutants for each—may further dysregulate intermediate progenitor proliferation resulting in altered transient divergence during peak L 2/3 PN genesis. These changes, however, are likely probabilistic based upon dynamic ranges of gene expression and diversity of cell cycle regulation across bP and NB subsets we have found. Reduced probability of aberrant cell states susceptible to replication stress due to 22q11 deletion—via diminished dosage of individual 22q11 genes, divergent epigentic regulation, or altered chromatin interactions reflecting the deletion itself^80, 81^—may restore typical L 2/3 PN genesis as proliferative activity and upper layer neurogenesis slow during late gestation.

Recent observations of constrained relationships between cortical progenitor identities and PN fates suggest that terminally dividing bPs yield daughter cells with similar laminar positions and cortico-cortical or subcortical projections^82^. Transient disruption of rapidly dividing bPs and their immediate NB progeny during peak upper layer neurogenesis may therefore amplify frequency of divergent L2/3 PNs or their vulnerability to subsequent pathogenic changes that result in aberrant association cortico-cortical connectivity. Additional consequences of mutations, CNVs, and environmental stressors may provide additional “hits” to further enhance pathology and dysfunction ^83, 84^. Nevertheless, changes due to disrupted L 2/3 PN progenitor cell states during a prolonged peak period of neurogenesis provide a likely foundation for subsequent cortical circuit pathology due to 22q11 deletion or additional mutations associated with elevated risk for schizophrenia and other neurodevelopmental disorders.

## ACKNOWLEDGMENTS

We thank Jill Maynard, Nate Faulkner, De’Onna Battle, Grace Boyer and Hannah Young for assistance in data collection and analysis. *Tbr2*^eGFP^ mice were obtained via MMRRC. Funding was provided by NICHD R01 HD042182 to ASL, the Seale Innovation Fund to TMM, and an award from the Red Gates Foundation to ASL.

