## Supplemental Figure Legends for "Transient disruption of progenitor states and layer 2/3 projection neuron generation in frontal association cortex due to 22q11.2 gene deletion"

**Supplemental Figure 1:** *Distribution of embryonic cortical precursors, neuroblasts and recently generated projection neurons.* Proportional distribution comparison of WT (white bars) and *LgDel* (black bars) * aPs, bPs, NBs, and ePNs across 20 proportionally equivalent bins (based upon cortical thickness measures in each animal analyzed from apical (ventricular) to basal (pial) surface in mFAC at E13.5 (***top row***), 14.5 (***second row***), 16.5 (***third row***) and E15.5 and 16.5 ePNs (***bottom row***). Asterisks indicate per-bin difference p<0.05(2-way ANOVA with Holm-Šídák's multiple comparisons test).

**Supplemental Figure 2:** *Similar transcriptional identities for WT and LgDel progenitor and NB subtypes.* ***top.*** Confirmation of identity and quantitative stability of additional aP, bP and NB markers across shared WT and *LgDel* aP, bP and NB subsets. ***bottom.*** Canonical upper and lower layer PN markers are expressed at similar levels in WT and *LgDel* aP, bP and NB subsets. Lighter half-circles denote WT, darker half-circles denote *LgDel*; frequency scale at bottom.

**Supplemental Figure 3.** *Expression of 22q11-deleted genes in WT and LgDel progenitor and NB subtypes***.** Expression of 22 22q11-deleted genes across all aP, bP, and NB subsets. Smaller dark half-circles indicate reduced mean expression in *LgDel* cells. Transcripts in gray are detected below the significance threshold applied to the scRNAseq data (mean ≥0.1 transcripts per cell).

**Supplemental Figure 4:** *Gene Set Enrichment Analysis (GSEA) of RNAseq data from Tbr2$^{eGFP+}$ cells.* GSEA of the bulk RNA sequencing dataset. ***left.*** Two-side barplot showing hallmark gene sets with significant enrichment (adjusted p < 0.05). Gene sets upregulated in *LgDel* bPs with positive normalized enrichment score (NES) are shown on the top right, while downregulated gene sets with negative (NES) are on the bottom left. ***middle***. Dot plot summarizing enriched pathways identified by GSEA. Each dot corresponds to a gene set or pathway. The x-axis shows the normalized enrichment score (NES), where positive values indicate upregulated/enriched gene sets and negative values indicate suppressed/underrepresented sets. Dot size reflects the number of overlapping genes from the dataset with each gene set, and dot color indicates statistical significance (adjusted p-value). ***right***. Individual GSEA enrichment plots are shown for five representative hallmark pathways with significant enrichment, illustrating the ranked gene list metric and the running enrichment score for each gene set.

**Supplemental Figure 5:** *Validation of 22q11 and DE genes in the E14.5 VZ/SVZ/IZ and CP.*

***top row***. Ventricular zone (VZ), subventricular zone (SVZ), intermediate zone (IZ), cortical plate (CP) and marginal zone (MZ) expression of DE genes in Tbr2eGFP+cells based upon colorimetric in situ hybridization in midline sagittal sections from the Genepaint (https://genepaint.org70) E14.5 fetal mouse dataset. Dark to light shading represents visually assessed labeling intensity in each cortical zone from ventricle to pial surface. All genes are localized to at least one developing cortical zone where Tbr2eGFP+ cells are found; asterisks indicate genes with rostral-caudal expression gradients. ***second-row***. Five DE genes expressed in anterior low-posterior high (*Slc26a1*, *Bcan*), anterior high-posterior low (*Gfra1*), anterior low-middle high-posterior low (*Meg3*) or anterior high-middle low-posterior high (*Col5a2*) gradients. ***third-row***. 22q11 gene expression in the E14.5 VZ, SVZ, IZ, CP and MZ based upon GenePaint *in situ* localization of 16 22q11 genes: asterisks indicate genes with rostro-caudal expression gradients. ***Third row, right***. ISH images from GenePaint were scored qualitatively based on visual signal intensity as not detected, very low, low, or high expression. Anterior low – posterior high gradients of Dgcr14, Gsc2 (excluding high olfactory bulb), and Slc25a1. ***bottom row.*** *Cdc45*, *Col5a2*, *Dlk1*, *Rian*, *Meg3*, *Ranbp1*, and *Bcan* expression levels quantified by qPCR in RNA samples from pooled fluorescence-activated cell sorted (FACS)-isolated *Tbr2*$^{eGFP+}$ cells from WT (black) and *LgDel* (magenta) samples parallel to those used for bulk RNA sequencing (n=6 WT, 6 *LgDel*, from 25 WT and 26 *LgDel* E14.5 fetuses). Results are presented as ΔCT values normalized to GAPDH; lower ΔCT indicates higher gene expression. Each data point represents a biological replicate corresponding to a single pooled sample. Individual data points reflect average values of three independent qPCR runs to ensure robustness. Statistical significance between WT and *LgDel* groups assessed using a paired two-tailed Student's t-test to account for batch effect.

**Supplemental Figure 6:** *scRNAseq confirmation of bulk RNA-seq identified DE genes across precursor and neuroblast subsets.* ***top***. Each bubble represents DE gene expression within a specific subset; light half-circles denote WT and dark half-circles denote LgDel. ***bottom***. Expression of Dlk1 locus genes is shown across aP, bP, and NB subsets based on scRNA-seq analysis. The bubble plot scale for Dlk1 and Rtl1 is magnified 5X to account for detection below the >0.1 transcripts-per-cell cutoff. Consistent with bulk RNAseq, expression levels of Dlk1, Meg3, and Rian are apparently increased in scRNAseq-defined bP1 as well as NB5 and NB6 subsets.

**Supplemental Figure 7:** *Analysis of frequency and distribution relationships for 22q11 deleted gene transcripts and Dlk1 locus transcripts in WT vs. LgDel VZ/SVZ Tbr2*$^{+}$ *and Tbr2*$^{-}$ *cells.* ***top left***. Significantly altered distribution/variance of *LgDel Cdc45* RNAscope puncta in individual Tbr2$^{-}$ VZ/SVZ cells (Levene's and Brown-Forsythe's tests). *<u>inset</u>*: Significantly reduced (per cell/per animal) mean *Cdc45* puncta frequency in *LgDel* Tbr2$^{-}$ (DAPI$^{+}$) VZ/SVZ cells (unpaired t-test). ***bottom left***. Significantly altered distribution/variance of *LgDel Dgcr8* RNAscope puncta in individual Tbr2$^{-}$ VZ/SVZ cells (Levene's and Brown-Forsythe's tests). *<u>inset</u>*: Reduced (per cell/per animal) mean *Dgcr8* puncta frequency in *LgDel* DAPI+Tbr2$^-$ VZ/SVZ cells (unpaired t-test) is not statistically significant, possibly because *LgDel Dgcr8* expression, based upon bulk RNAseq and qPCR, is higher than the 50% average of other 22q11 genes. ***top middle***. *Meg3* puncta distribution/variance in individual *LgDel* Tbr2$^-$ VZ/SVZ cells is not altered (Levene's and Brown-Forsythe's tests). *inset*: Mean (per cell, per animal) WT vs. *LgDel Meg3* puncta frequency does not differ significantly (Unpaired t-test). ***bottom middle***. *Rian* puncta distribution/variance in individual *LgDel* Tbr2$^-$ VZ/SVZ cells is not altered (Levene's and Brown-Forsythe's tests). *inset*: Mean (per cell, per animal) WT vs. *LgDel Rian* puncta frequency does not differ significantly (Unpaired t-test). ***top middle.*** Very weak (WT) to weak (*LgDel*) correlation between *Cdc45* (*C45*) and *Dgcr8* (*D8*) puncta frequency per Tbr2$^+$ cell. ***bottom middle.*** Weak (*LgDel*) to very weak (WT) correlation between *Cdc45* and *Dgcr8* puncta frequency per Tbr2$^-$ cell. ***top right*.** Very weak correlation between *Dgcr8* puncta frequency and Tbr2$^+$ cell size (Tbr2$^+$ nuclear area) in WT and *LgDel*. ***bottom right.*** Very weak correlation between *Dgcr8* puncta frequency and Tbr2$^-$ cell size (DAPI$^+$ nuclear area) in WT and *LgDel*.

**Supplemental Figure 8:** *Spatiotemporal distribution of Cux1$^+$ and Cux1$^-$ cortical neurons in the P5 mFAC and V1.* ***top.*** Distribution comparison of Cux1$^+$ (red) and Cux1$^-$ (grey) across 20 proportionally equivalent bins spanning from the apical to basal surface in the medial frontal association cortex (mFAC) for WT (left) versus *LgDel* (right) pups birthdated across developmental stages E12.5, E13.5, E14.5, E15.5, and E16.5 (ordered left to right). ***bottom.*** Distribution comparison of Cux1$^+$ (red) and Cux1$^-$ (grey) in the primary visual cortex (V1) birthdated across E13.5 to E16.5 (ordered left to right), comparing WT (left) and *LgDel* P5 pups.

**Supplemental Figure 9:** *Analysis of birthdated projection neurons in the postnatal primary* *somatosensory cortex (S1).* ***left.*** Representative images of primary somatosensory cortex birthdate at E14.5 with IdU (green) and at E16.5 with BrdU (cyan); Brn2$^+$ cells (red) are seen in L 2/3 as well as lower L 4. ***middle***: Frequency of WT and *LgDel* E14.5 and E16.5 birthdated neurons assessed at P5 in primary somatosensory cortex (unpaired t-test, n=5 WT, 7 *LgDel*). ***right.*** Distribution analysis of WT (black) and *LgDel* (grey) E14.5 (left) and E16.5 (right) birthdated cells across 20 proportionally equivalent bins from the apical to basal surface.

**Supplemental Figure 10**: *Similar distribution and frequency of Brn2$^+$ L 2/3 PNs in WT and LgDel P5 mFAC.* ***left.*** Brn2$^+$ L 2/3 PNs show no significant differences between genotypes in frequency and laminar positions of this subset of upper layer PNs in mFAC. Y-axis divided into 20 proportional bins, apical to basal surface, based upon cortical probe thickness for each animal, ***inset***: Mean Brn2$^+$ PN frequency per animal (unpaired t-test, n=5 WT, 5 LgDel) confirms overall equivalence in the two genotypes**.** ***right.*** Parallel analysis of the V1 (right panel) similarly demonstrates equivalent Brn2$^+$ PN frequency and laminar distribution across both genotypes (unpaired t-test, n = 7 WT, 6 *LgDel*).
