## Supplementary figures and images for "Transient disruption of progenitor states and layer 2/3 projection neuron generation in frontal association cortex due to 22q11.2 gene deletion"

### Supplemental Figure 1

**Rukh et al. 2026 Supplemental Figure 1**

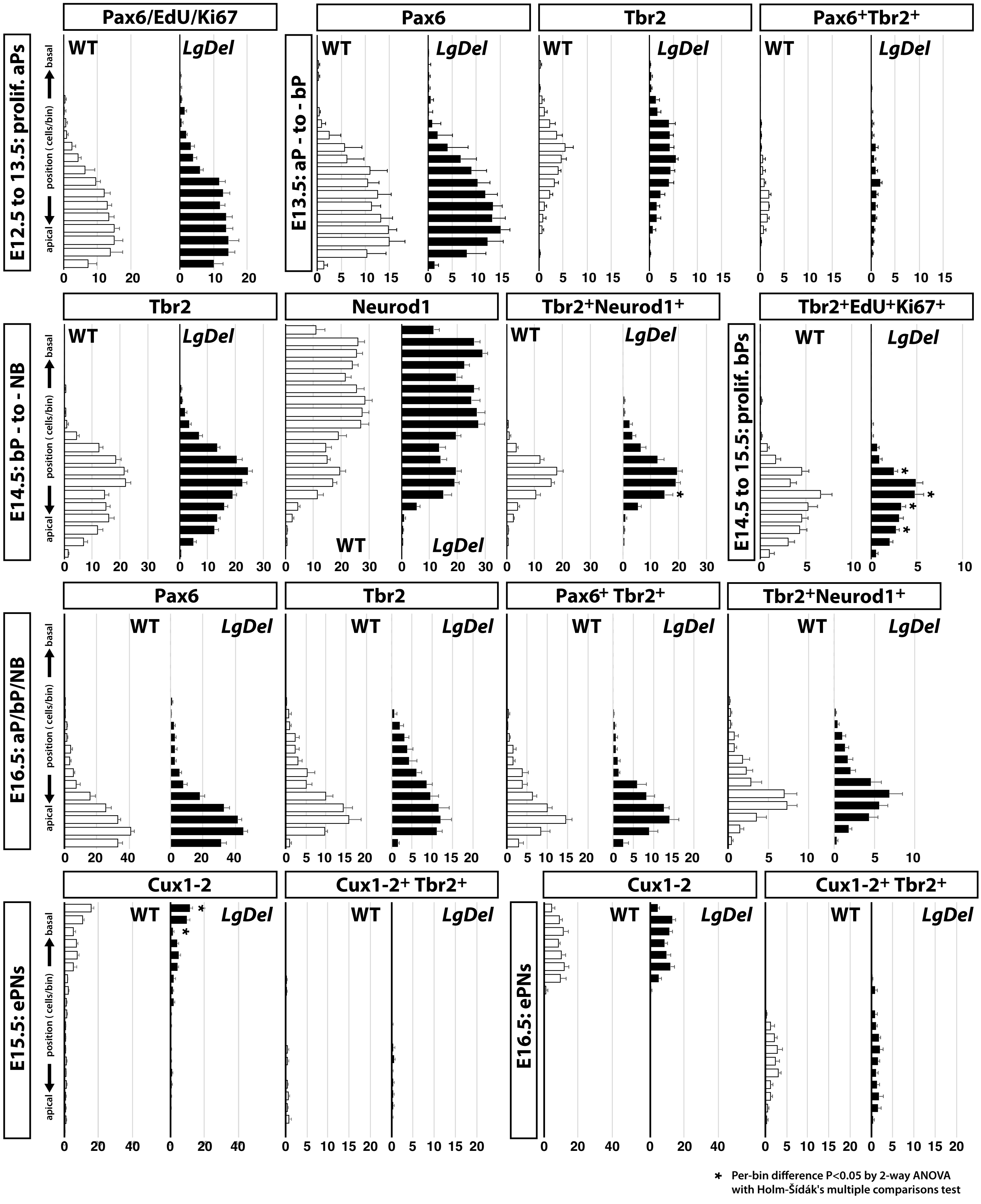

### Supplemental Figure 2

**Rukh et al. 2026** **Supplemental Figure 2**

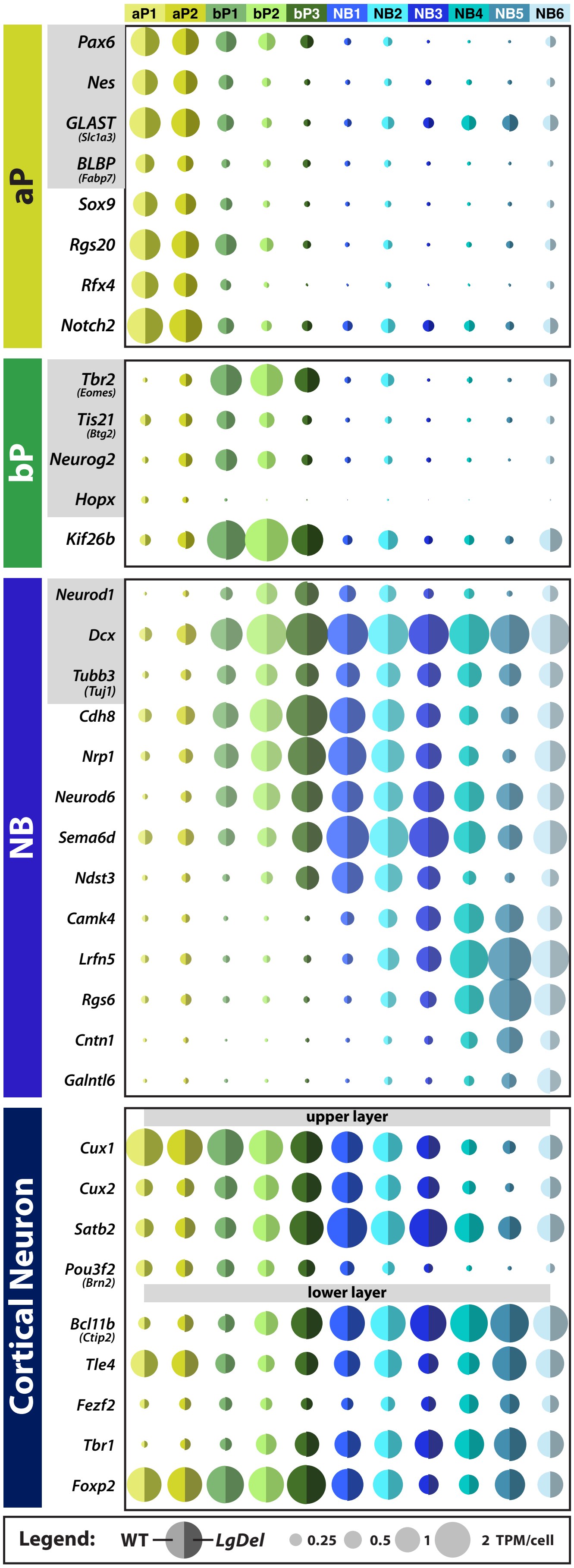

### Supplemental Figure 3

**Rukh et al. 2026** **Supplemental Figure 3**

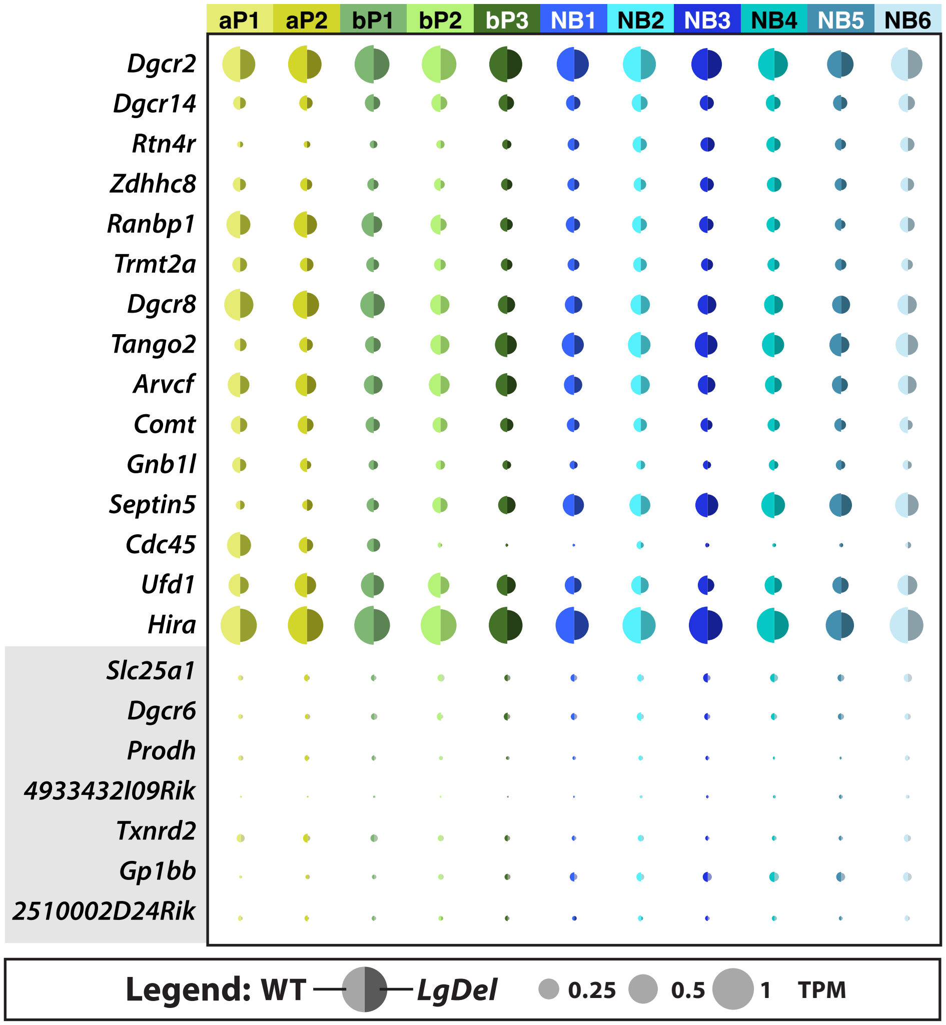

### Supplemental Figure 4

#### Rukh et al. 2026 Supplemental Figure 4

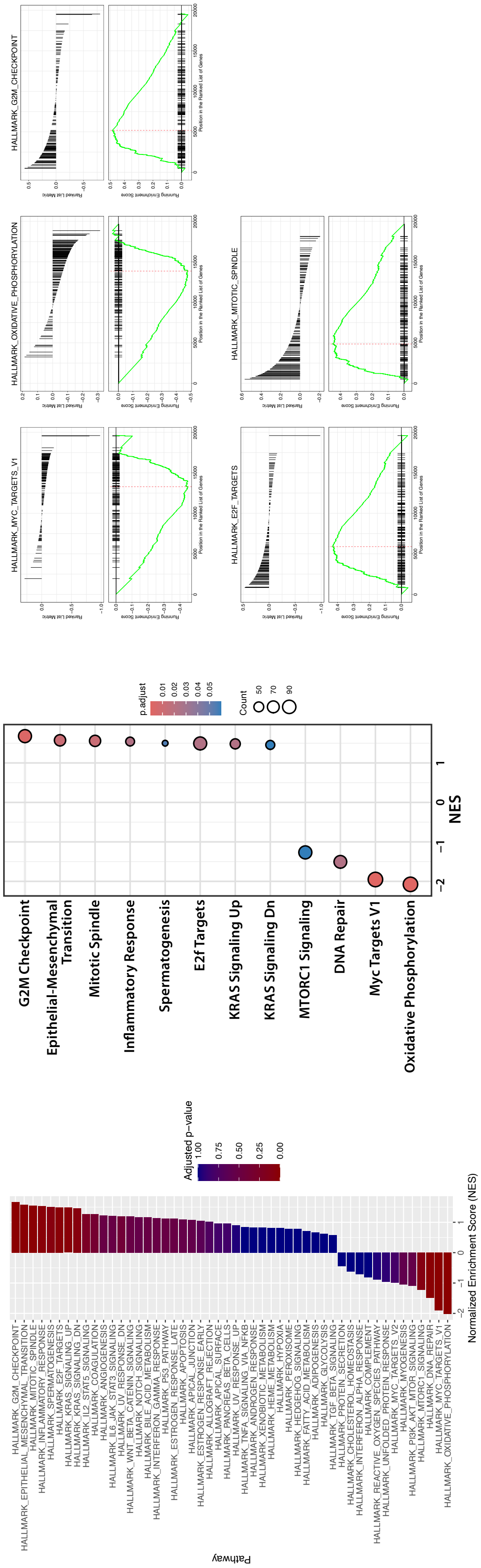

### Supplemental Figure 5

#### Rukh et al. 2026 Supplemental Figure 5

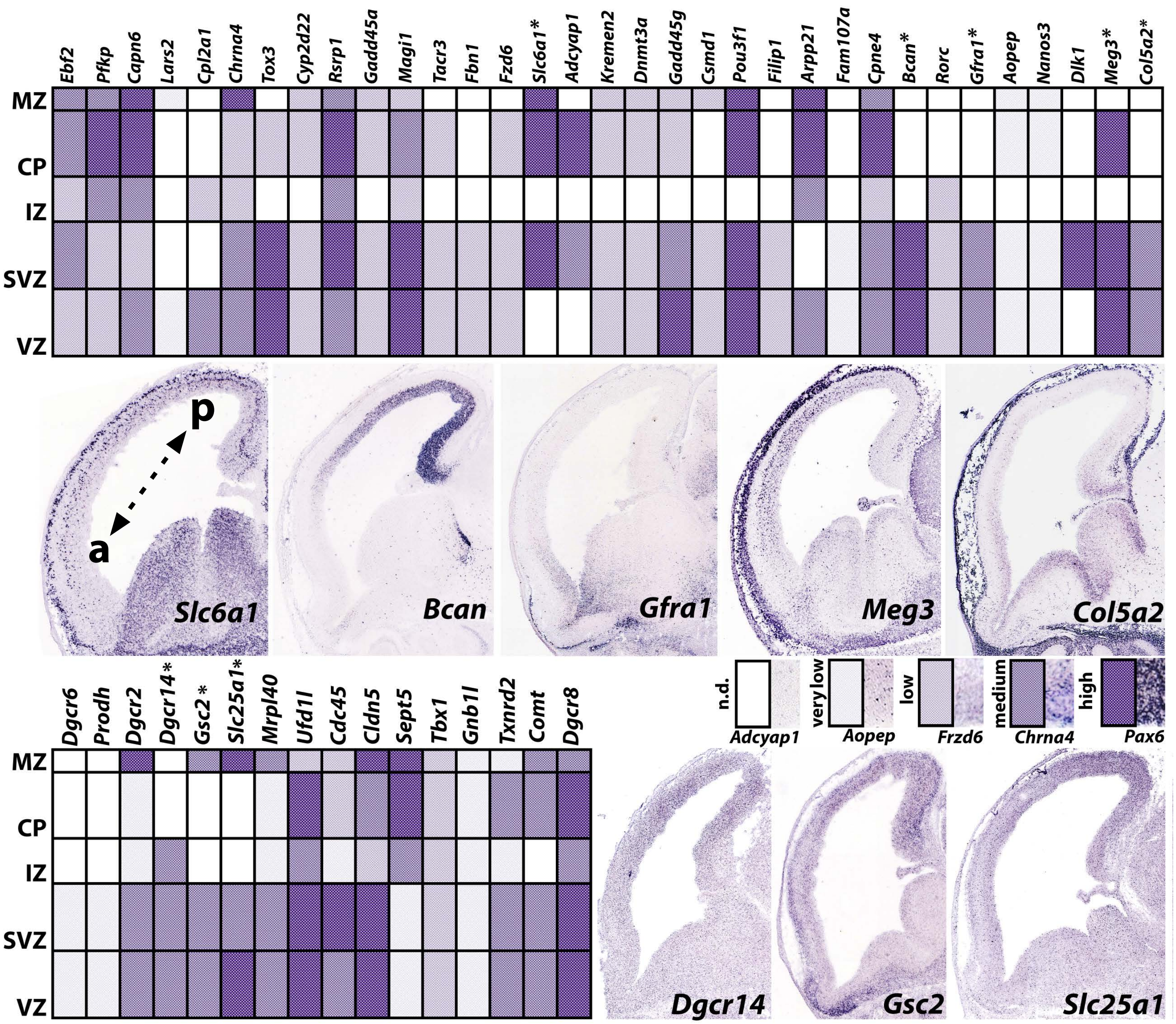


**qPCR validation of candidate genes from sorted BP samples**

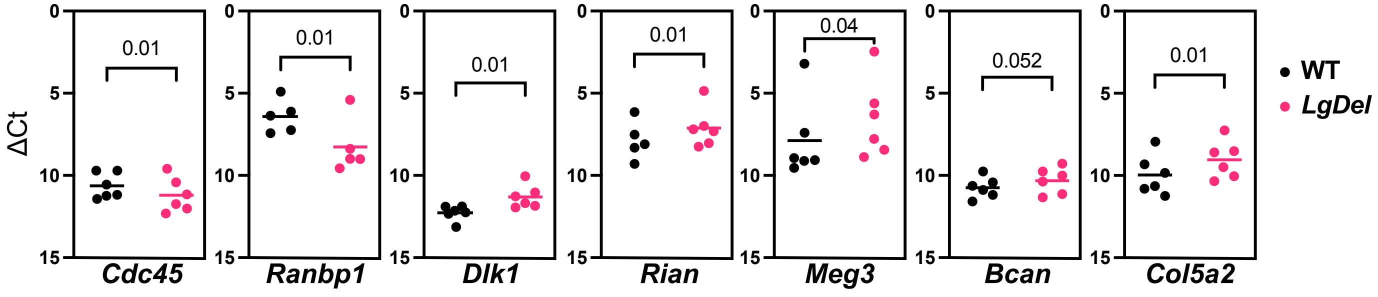

### Supplemental Figure 6

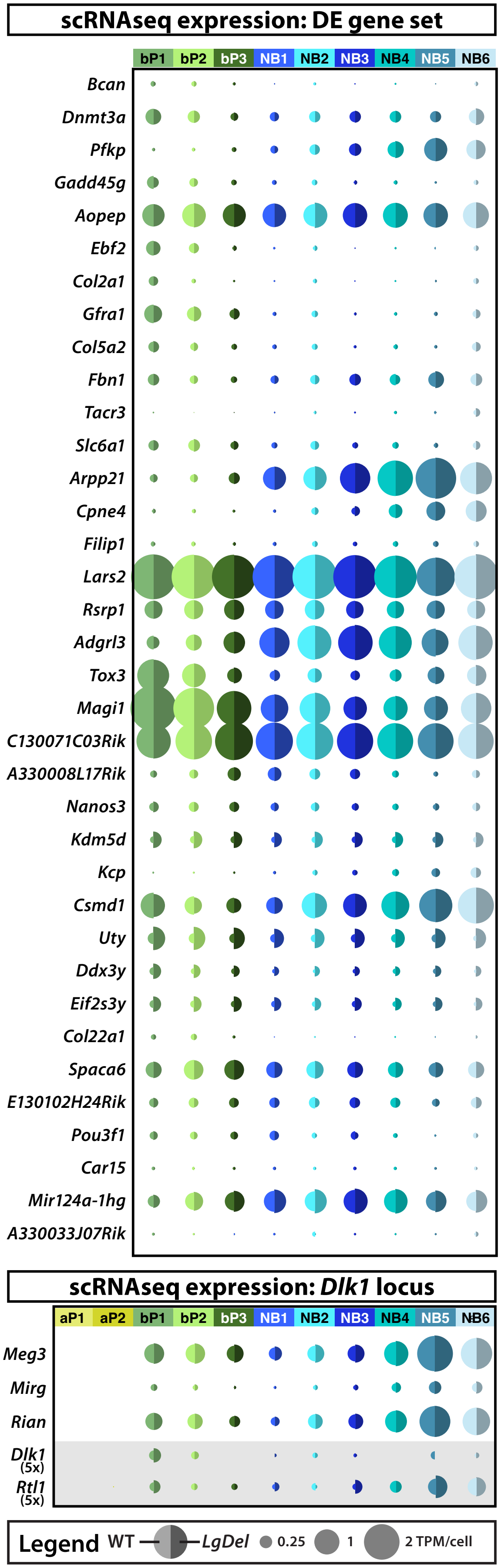

### Supplemental Figure 7

#### Rukh et al. 2026 Supplemental Figure 7

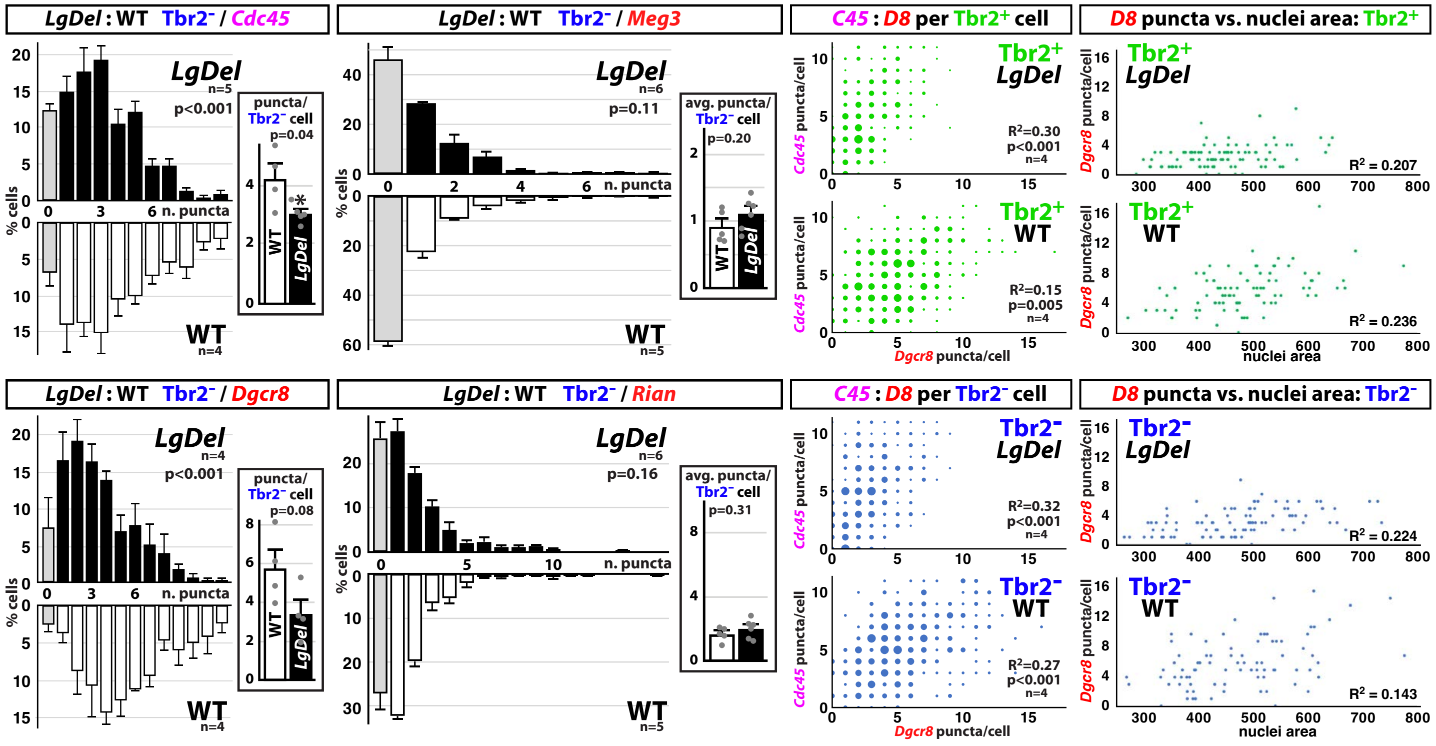

### Supplemental Figure 8

#### Rukh et al. 2026 Supplemental Figure 8

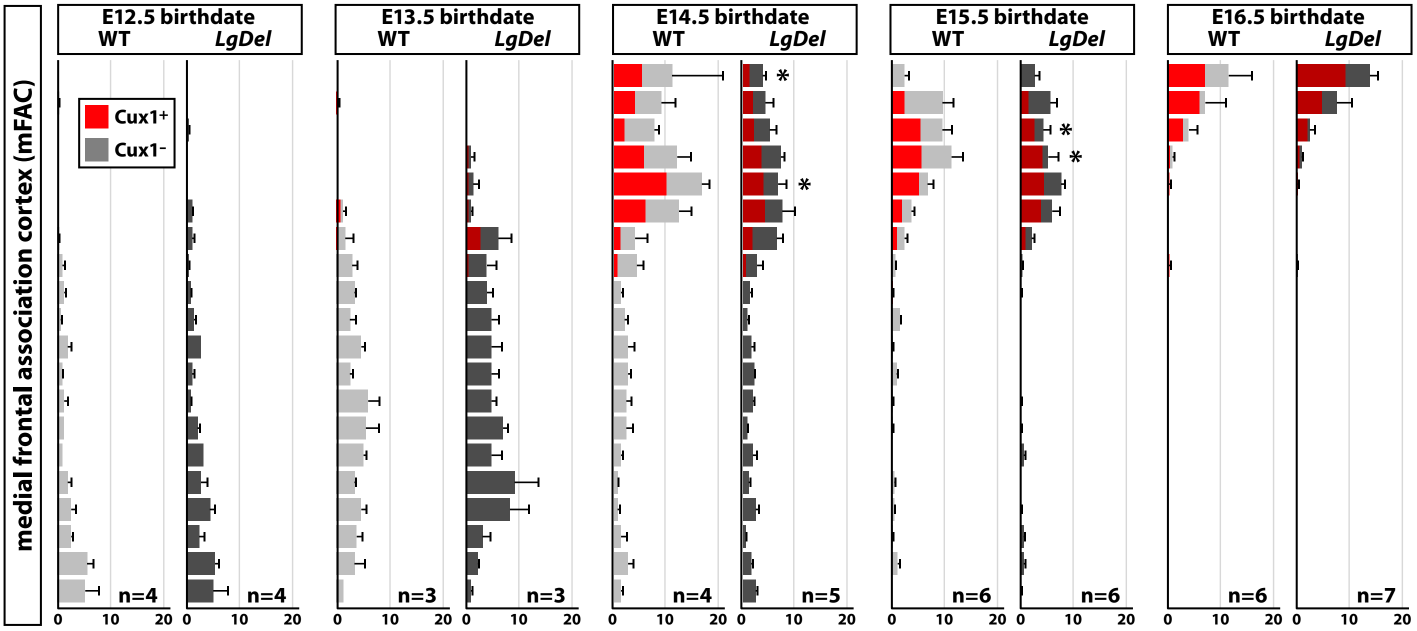


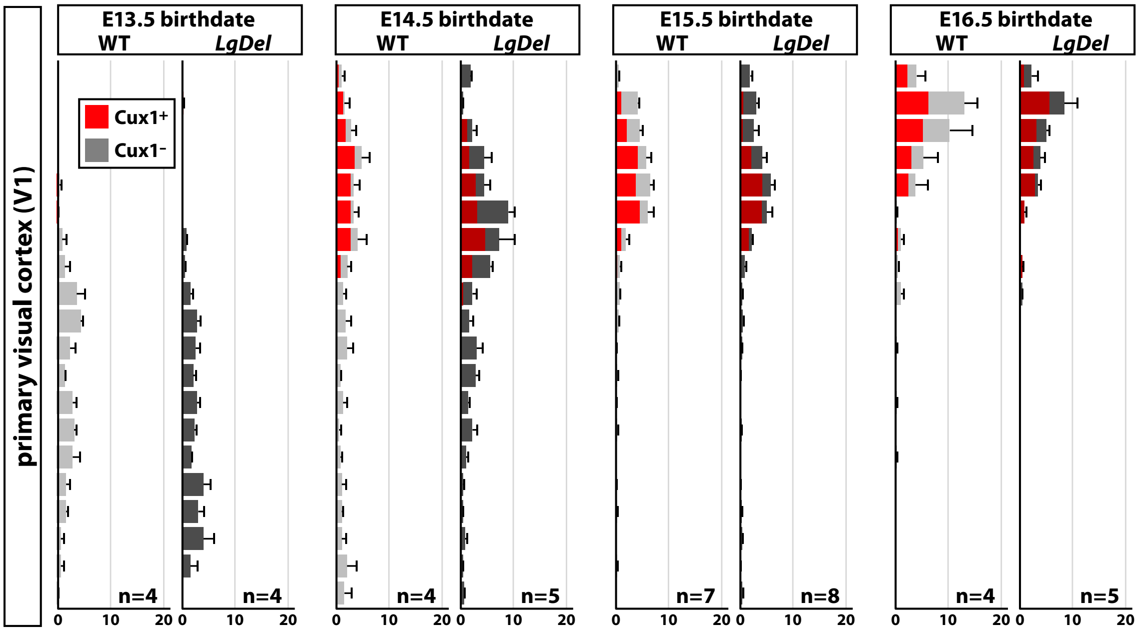

### Supplemental Figure 9

#### Rukh et al. 2026 Supplemental Figure 9

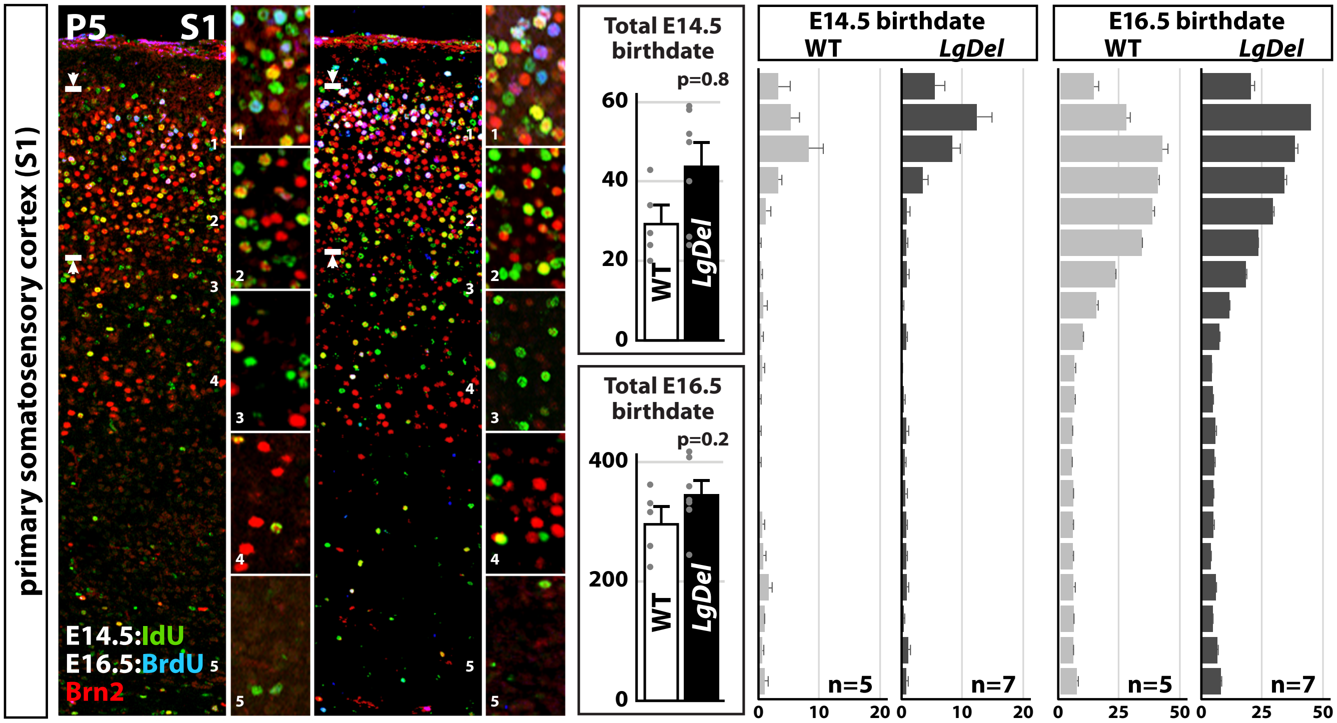
