## Supplemental Figure 10 for "Transient disruption of progenitor states and layer 2/3 projection neuron generation in frontal association cortex due to 22q11.2 gene deletion"

Rukh et al. 2026 Supplemental Figure 10

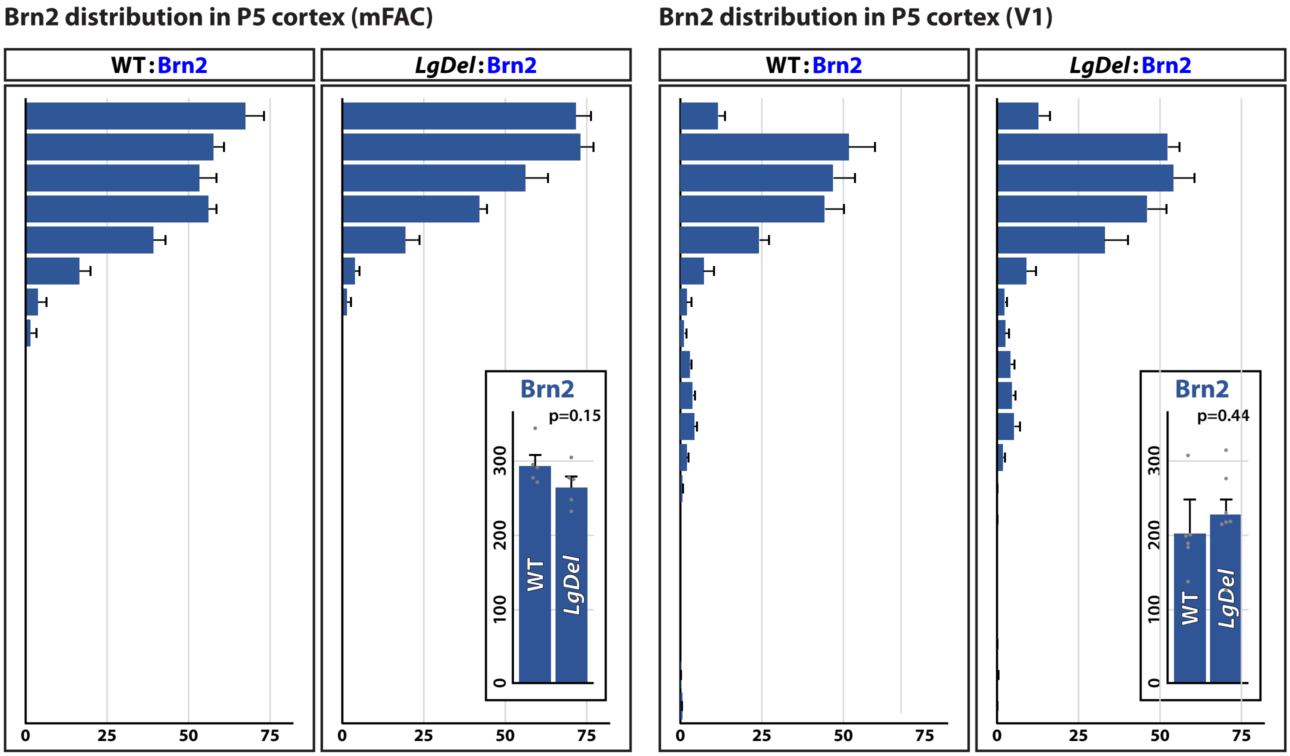


Distribution plot is parallel to Fig. 5b, c

Distribution plot is parallel to Fig. 5f, g
