## Supplemental Table Legends for "Transient disruption of progenitor states and layer 2/3 projection neuron generation in frontal association cortex due to 22q11.2 gene deletion"

**Supplemental Table 1**: PCR primers used for quantitative (qPCR) validation of differentially expressed genes (top) as well as genotyping of *LgDel* vs. WT fetuses and postnatal mice.

**Supplemental Table 2**: Primary (top) and secondary (bottom) antibodies used to identify progenitor and projection neuron cell types as well as S-phase labeled cells. The species and type is provided for each antibody, as well as source (with catalog number), concentration used, and antigen retrieval conditions, when used.

**Supplemental Table 3**: RNAscope probes used for cellular validation of 22q11 and differentially expressed gene expression levels. All probes from ACD/Biotechne; catalogue numbers and targeted base pairs are provided as available.

**Supplemental Table 4**: PCR primers for amplicons to tile the Meg3 differentially methylated region (DMR) as well as the *Dlk1* locus intergenic DMR (IG-DMR), two locus control regions whose methylation influences expression of 5 genes within the *Dlk1* locus.
