## Supplemental Table 1 for "Transient disruption of progenitor states and layer 2/3 projection neuron generation in frontal association cortex due to 22q11.2 gene deletion"

| qPCR primers | | |
|---|---|---|
| **Primer** | **Forward** | **Reverse** |
| Cdc45 | CAG AGA AGC GCA CAC GGT TAG AAG AG | CCC ATG TCT GCA AGG AAC TCC TGG AG |
| Ranbp1 | GAC CCC CAG TTC GAG CCA ATA GTT TC | CAT TTA GGA AGC GGA TGG CGA GCA G |
| Dlk1 | CCTGGCTGTGTCAATGGAGT | TGGCAGTCCTTTCCAGAGAA |
| Meg3 | CGAGGACTTCACGCACAA | ATTCCAGATGATGGCTTTG |
| Rian | TGTCGGAGGATCGTGTCAT | AATCTTCTAGAGCCCAGATCC |
| Col5a2 | GGAGAAGGAAACATCAGATTCA | CCAAATTCCTGGTCTGCATT |
| Bcan1 | CTATGTTTGCCAGGCTATGGGGG | TGCCTCCTCCCAACTCCTTCGTG |
| Gapdh | CTG ACG TGC CGC CTG GAG AAA | GTT GGG GGC CGA GTT GGG ATA GG |
| **genotyping primers** | | |
| **Primer** | **Forward** | **Reverse** |
| LgDel | GGCCAGCTCATTCCTCCCACTC | CACCAATCACAGGGCTGGAGGACTAC |
| C45 | GGCCTCGGACGTGGATGCCTTGTG | CGGGGCTCCTGTATTTCCTGCACTC |
