## Supplemental Table 2 for "Transient disruption of progenitor states and layer 2/3 projection neuron generation in frontal association cortex due to 22q11.2 gene deletion"

| | Antibody Name | Clonality Species | Source | Catalog No. | Concentration | Antigen Retrieval |
|---|---|---|---|---|---|---|
| Primary Antibodies | Tbr2 | guinea pig polyclonal antibody | Synaptic Systems | 483005 | IHC (1:1000) | Treated with sodium citrate pH 7.0 at 95°C for 10 min |
| | Satb2 | guinea pig polyclonal antibody | Synaptic Systems | 327004 | IHC (1:500) | Treated with sodium citrate pH 7.0 at 95°C for 10 min |
| | BrdU (+ IdU) | chicken polyclonal antibody | Innovative research | 43421 | IHC (1:500) | Treated with sodium citrate pH 7.0 at 95°C for 15 min |
| | BrdU | rat monoclonal antibody | Abcam | Ab6326 | IHC (1:100) | Treated with sodium citrate pH 7.0 at 95°C for 15 min |
| | Brn2 | mouse monoclonal antibody | Santa Cruz Biotechnology | 393324 | IHC (1:100) | Treated with sodium citrate pH 7.0 at 95°C for 10 min |
| | Pax6 | rabbit polycolnal antibody | Biolegend | 9013901 | IHC & Pair cell assay (1:1000) | IHC: Treated with sodium citrate pH 7.0 at 95°C for 10 min |
| | Tbr2 | rabbit polycolnal antibody | Abcam | Ab23345 | IHC & Pair cell assay (1:1000) | Not required |
| | Neurod1 | mouse monoclonal antibody | Abcam | Ab60704-1001 | IHC (1:350) | Required RNAScope manual assay protocol |
| | Tbr2 | chicken polycolnal antibody | Millipore | A615894 | IHC (1:200) | Treated with sodium citrate pH 7.0 at 95°C for 10 min |
| | Ki67 | mouse monoclonal antibody | BD pharminogen | 556003 | IHC (1:400) | Treated with sodium citrate pH 7.0 at 95°C for 10 min |
| | Ki67 | rat monoclonal antibody | Biolegend | 652402 | IHC (1:200) | Treated with sodium citrate pH 7.0 at 95°C for 10 min |
| | Cux1/2 | rabbit monoclonal antibody | Abcam | Ab309139 | IHC (1:50) | Treated with sodium citrate pH 7.0 at 95°C for 10 min |
| | Cux1 | rabbit polycolnal antibody | Proteintech | 11733-1-AP | IHC (1:1000) | Treated with sodium citrate pH 7.0 at 95°C for 10 min |
| Secondary Antibodies | Alexa Fluor 488 anti-Chicken | Goat polyclonal antibody | Invitrogen | A11039 | IHC (1:4000) | 1-hour incubation at RT |
| | Alexa Fluor 546 goat anti-Chicken | Goat polyclonal antibody | Invitrogen | A11040 | IHC (1:2000) | 1-hour incubation at RT |
| | Alexa Fluor 647 goat anti-Chicken | Goat polyclonal antibody | Invitrogen | A32933 | IHC & Pair cell assay (1:2000) | 1-hour incubation at RT |
| | Alexa Fluor 633 goat anti-Rat | Goat polyclonal antibody | Invitrogen | A21094 | IHC (1:2000) | 1-hour incubation at RT |
| | Alexa Fluor 647 goat anti-Rat | Goat polyclonal antibody | Invitrogen | A21248 | IHC (1:2000) | 1-hour incubation at RT |
| | Alexa Fluor 488 goat anti-Rabbit | Goat polyclonal antibody | Invitrogen | A11034 | IHC (1:4000) | 1-hour incubation at RT |
| | Alexa Fluor plus 405 goat anti-Rabbit | Goat polyclonal antibody | Invitrogen | A48254 | IHC (1:500) | 1-hour incubation at RT |
| | Alexa Fluor 647 goat anti-Rabbit | Goat polyclonal antibody | Invitrogen | A21244 | IHC (1:2000) | 1-hour incubation at RT |
| | Alexa Fluor 546 goat anti-Rabbit | Goat polyclonal antibody | Invitrogen | A11035 | IHC (1:2000) | 1-hour incubation at RT |
| | Alexa Fluor 680 goat anti-Mouse | Goat polyclonal antibody | Invitrogen | A21057 | IHC (1:2000) | 1-hour incubation at RT |
| | Alexa Fluor 546 goat anti-Mouse | Goat polyclonal antibody | Invitrogen | A11030 | IHC & Pair cell assay (1:2000) | 1-hour incubation at RT |
| | Alexa Fluor 647 goat anti-Mouse | Goat polyclonal antibody | Invitrogen | A21235 | IHC (1:2000) | 1-hour incubation at RT |
| | Alexa Fluor 546 goat anti-Guinea pig | Goat polyclonal antibody | Invitrogen | A11074 | IHC (1:2000) | 1-hour incubation at RT |
| | Alexa Fluor 647 goat anti-Guinea pig | Goat polyclonal antibody | Invitrogen | A21450 | IHC (1:2000) | 1-hour incubation at RT |
