## Supplemental Table 3 for "Transient disruption of progenitor states and layer 2/3 projection neuron generation in frontal association cortex due to 22q11.2 gene deletion"

| Gene Name | Catalog No. | Target Region Base Pairs (bp) | RNAscope Assay Platform |
|---|---|---|---|
| Dlk1 | 405971-C3 | 1477 - 2387 | Manual Assay RNAscope |
| Meg3 | 527201 | 2043 - 3118 | Manual Assay RNAscope |
| Rian | 510531 | 388 - 1387 | Manual Assay RNAscope |
| Dgcr8 | 1243221-C3 | 664 - 1667 | Manual Assay RNAscope |
| Cdc45 | 832311 | 129-1042 | Manual Assay RNAscope |
