## Supplemental Table 4 for "Transient disruption of progenitor states and layer 2/3 projection neuron generation in frontal association cortex due to 22q11.2 gene deletion"

| Bisulphite specific PCR primers | | | | | | |
|---|---|---|---|---|---|---|
| Name of the target | Forward Primer Sequence | Reverse Primer Sequence | Start with primer | End with primer | Chromosome | Strand |
| Meg3_DRM_1 | AYGTTTTTATTTTTGAAAAGGTTTGAGTTATTT | CCCCCTATAACCAACAAACCTAAAATAC | 109539619 | 109539764 | chr12 | + |
| Meg3_DRM_2 | GGTTTTTTTAGTTTTATGTTTTTTATAGGGTTTTTG | CCRACTACTCTACCTACCCTAAATAA | 109539824 | 109540055 | chr12 | + |
| Meg3_DRM_3 | GAGAGGATATTTGATGGATATGTTGTTTTTT | CCCTATAATAATACTTAAAATACACCCCCAA | 109539995 | 109540228 | chr12 | + |
| Meg3_DRM_4 | YGTTTTTTAAAGTGTGGGGAATTAGTT | AAAACCTTAATTTTAAATCCCAAACTTTACAAA | 109540166 | 109540417 | chr12 | + |
| Meg3_DRM_5 | GGTGTATAGGTTGTATTTTTTGTAAAGTTTGG | CTAAATTTCTCAAAACATCAATCAAAAACTCC | 109540366 | 109540623 | chr12 | + |
| Meg3_DRM_6 | GTTTTTGATTGATGTTTTGAGAAATTTAGGTAAG | CCAAATTCTATAACAAATTACTATAACCRCAC | 109540594 | 109540878 | chr12 | + |
| Meg3_DRM_7 | GTGYGGTTATAGTAATTTGTTATAGAATTTGG | CAAAAAATAACTAACCACTCACCAAATAAAAAATTC | 109540846 | 109541121 | chr12 | + |
| Meg3_DRM_8 | GGTTAGTTATTTTTTGTTTGAAAGGATGTG | CTCATTATTACTACTTTTCTTAATCAAAATCTCTACC | 109541105 | 109541392 | chr12 | + |
| Meg3_DRM_9 | GGTTTTGGTGGTTGAAAGTTTTTTTTAG | CAAAATTACTAATCAACATAAACCTCTAAATCTTAC | 109541443 | 109541640 | chr12 | + |
| Meg3_DRM_10 | GTAAGATTTAGAGGTTTATGTTGATTAGTAATTTTG | TAAAATTTTTTCTACCCTAACCCACCC | 109541604 | 109541866 | chr12 | + |
| Meg3_DRM_11 | GAGAGGGYGTTAAAGGTAAATTTAGT | CTCCAAATTTCCAACACTCAAATCAC | 109541750 | 109541958 | chr12 | + |
| Meg3_DRM_12 | GTGATTTGAGTGTTGGAAATTTGGAG | CCATCAATAAAACATTCTAAAAACCTACCA | 109541932 | 109542169 | chr12 | + |
| IG_DMR_1 | GTGTTTTAGGTTTGTGTTGTGAGGTATTA | ACCATAATATACTACTCCTCAAAACCCA | 109526409 | 109526587 | chr12 | + |
| IG_DMR_2 | GGGTTTTGAGGAGTAGTATATTATGGT | ACTATTATATCACACTTAACTTTACTACTACACATC | 109526560 | 109526773 | chr12 | + |
| IG_DMR_3 | GTAAAGTTAAGTGTGATATAATAGTTTTGATTTGGT | CATAAAAACATTACTCCATAATACATCCTATTCC | 109526748 | 109527009 | chr12 | + |
| IG_DMR_4 | GGAATAGGATGTATTATGGAGTAATGTTTTTATG | CCACATATCTATACTACAAAACACCAAAC | 109526975 | 109527188 | chr12 | + |
| IG_DMR_4-1 | GTGGGTGTATTATGTATGGTATAAGATAGATG | ACACCTAACCTATACTACAAAACAATCA | 109527184 | 109527350 | chr12 | + |
| IG_DMR_5 | GGTATAAGATTGAATGTTTTTGTGGTAAAGG | CATCCATATATACCAAACCAATACTATAATACAC | 109527287 | 109527451 | chr12 | + |
| IG_DMR_6 | GGTATATATGGATGTATTGTAATATAGGTTAGGTG | CCATACACTTTTACTATAAAACACTTAACTC | 109527437 | 109527712 | chr12 | + |
| IG_DMR_7 | GGTATATTTTGGTATAGATTTGGTGTTTTATGG | TATACCATAACACAACTACACAAAATACTACA | 109527752 | 109527959 | chr12 | + |
| IG_DMR_8 | GGTGTATTTTGTATTATTGTTTTGTGTTAGAGTAG | CCTTCCTAACACTAACATAATATACCTTAACAC | 109527851 | 109528120 | chr12 | + |
| IG_DMR_8-1 | GTGTTAAGGTATATTATGTTAGTGTTAGGAAGG | CATAAAACTTACCCATAACAAACCACAA | 109528087 | 109528231 | chr12 | + |
| IG_DMR_9 | GTTAYGGTTTATAGTGGATAGTTAGTGT | TTCTCCATTAACAAAATAATACAACCCTTC | 109528270 | 109528547 | chr12 | + |
| IG_DMR_10 | TTTGTTAATGGAGAATGTTTTGAGTATAGG | AAACTCTACAAATACCAAATTCCATCAAA | 109528532 | 109528694 | chr12 | + |
| IG_DMR_11 | GGTAGTTTGTGAGTTGATTTTTTTTAGTTATAGT | CTATATAATCCACAATCTAACAAAACTACAACAAA | 109528781 | 109528924 | chr12 | + |
